# Novel Integrative Modeling of Molecules and Morphology across Evolutionary Timescales

**DOI:** 10.1101/242875

**Authors:** Huw A. Ogilvie, Fábio K. Mendes, Timothy G. Vaughan, Nicholas J. Matzke, Tanja Stadler, David Welch, Alexei J. Drummond

**Author notes:** Correspondence to be sent to: Huw A. Ogilvie, Department of Computer Science – MS-132, Rice University, P.O. Box 1892, Houston, TX 77251-1892, USA. Correspondence to be sent to: Fábio K. Mendes, School of Biological Sciences, The University of Auckland, Private Bag 92019, Auckland Mail Centre, Auckland 1142, New Zealand. These authors contributed equally (Huw A. Ogilvie, Fábio K. Mendes).

## Abstract

Evolutionary models account for either population- or species-level processes, but usually not both. We introduce a new model, the FBD-MSC, which makes it possible for the first time to integrate both the genealogical and fossilization phenomena, by means of the multispecies coalescent (MSC) and the fossilized birth-death (FBD) processes. Using this model, we reconstruct the phylogeny representing all extant and many fossil Caninae, recovering both the relative and absolute time of speciation events. We quantify known inaccuracy issues with divergence time estimates using the popular strategy of concatenating molecular alignments, and show that the FBD-MSC solves them. Our new integrative method and empirical results advance the paradigm and practice of probabilistic total evidence analyses in evolutionary biology.

## Introduction

Creating a high-resolution picture of the tree of life is an increasingly achievable goal given the ever greater availability of molecular and paleontological data. Realistic and tractable evolutionary models are required to treat this rich data in a statistically sound manner. The end result should be phylogenies that not only explain how species are related, but are also scaled to absolute time, which allows species trees to be reconciled with geological and fossil records.

One method for scaling trees into absolute time is to assume a molecular clock [1] ticking at a known rate (or rates) per unit time. This strategy is problematic because a universal clock does not exist, and extrapolating clock rates measured in one group of organisms to another can lead to unrealistic evolutionary time estimates [2, 3].

Alternatively, the “node dating” method [4, 5] proposes prior distributions for divergence times based on fossil ages and morphology. Yet this method faces many problems. Node dating only uses the oldest available fossils, ignoring younger fossils. Fossil affinities and associated node age priors are ultimately specified using expert knowledge [6] which, due to its ad hoc nature, can introduce explicit bias and circularity to divergence time estimates [7, 8]. Finally, the interaction between priors on “dated” nodes and the overall tree prior used in a hierarchical models creates complex and unintended prior probabilities on node ages throughout the tree [9].

The fossilized birth-death (FBD) model introduced probabilistic “tip dating” to paleontology and systematics [4, 10–12]. This model not only directly solves the shortcomings of node dating, but in providing more accurate model-based uncertainty on divergence times, it also allows relaxed clock models to be less distorted by inadequacies in the tree prior and calibration scheme. Unless fossil ages are data, relaxed clock models by themselves do not “close the gap between rocks and clocks” [13, 14], and only tell us about relative differences in accumulated evolutionary change, where time and rates are conflated. By using the FBD model, one can combine morphological characters and fossil ages with molecular data in a statistically robust framework, and disentangle absolute time from evolutionary rates.

Studies employing the FBD model have invariably assumed that morphological and molecular characters evolve along the same phylogeny. This assumption is the core of “concatenation” (initially referred to as “total evidence” data combination [15, 16]), a protocol that attempts to harness as much information from as many different data sources as possible. The hope of concatenation is that agreeing signals speak louder than the sampling noise, and that conflicting signals can compete in the resolution of the phylogenetic estimate. The crucial feature of concatenation, as opposed to integrative probabilistic models (discussed below), is that all characters are simply appended together into a single large data matrix.

Since genomes have become central data sources for studying the evolution of living species, concatenation is now often taken to mean “pasting” all sequenced nucleotides together into a single multiple sequence alignment (MSA). This is the meaning we employ here. Within the domain of molecular phylogenetics, concatenation has been shown to produce biased tree estimates in a maximum-likelihood context [17, 18]. In a Bayesian context, concatenation has been associated with the overestimation of tip branch lengths by as much as 350%, as well as inaccurately narrow credible intervals, which often exclude true parameter values and tree topologies [19–21]. By contrast, the multispecies coalescent (MSC) accurately models the evolution of multiple unlinked loci. Concatenation is still used due to the perceived higher computational cost of MSC, which we will show does not exist (relative to Bayesian concatenation) when inferring species trees using tip dating on a real data set.

Under the MSC, incomplete lineage sorting (ILS) can occur, where gene lineages fail to coalesce in their immediately common ancestral populations. In such events, depending on how lineages then sort, gene tree topologies might differ from the species tree topology [22, 23]. The MSC is demonstrably more accurate than concatenation when estimating topologies and relative branch lengths in simulated uncalibrated scenarios [20, 21], but has not yet been put to test with empirical data sets comprised of both multiple unlinked loci as well as morphological data.

We propose integrative models for species- and population-level evolution, as well as for speciation, extinction and fossilization processes, in order to leverage data of different kinds – molecules, morphology, and the fossil record – in a single probabilistic “total-evidence” [4] analysis. Our model circumvents the known issues caused by concatenation, while explicitly distinguishing the evolutionary processes behind species branching patterns and fossilization, and those behind genealogical branching patterns (Fig. 1). We call our new combined model FBD-MSC, implement it in StarBEAST2 [21] for the BEAST 2 platform [24], demonstrate its correctness, and then compile an exemplar data set of the Caninae (a major canid subfamily) with which we show our model in use.

**Figure 1:**
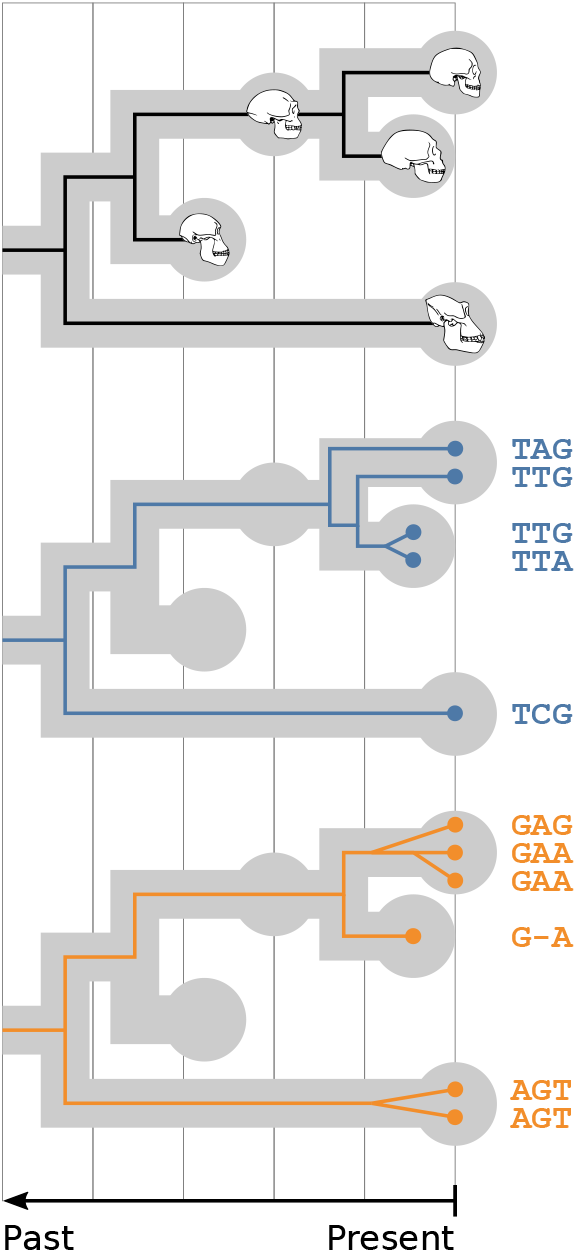
A species tree with a single sampled ancestor—a direct ancestor of other species in the tree—and its relationship to morphological data (top) and multilocus sequence alignments (middle and bottom) in a unified model. The fossilized birth-death (FBD) process is used to model fossilization, speciation and extinction processes, while the multispecies coalescent (MSC) is used to model gene tree evolution within the species tree. GTR family substitution models may be used to model sequence and trait evolution along gene and species trees respectively.

### Integrative Model Probability

The integrative model combining the MSC, the FBD process, and morphological evolution can be expressed by combining the probability mass and density functions (pmf and pdf) characterizing all the component sampling distributions. The probability of the *i*-th gene tree given the multiple sequence alignment (MSA) of the *i*-th gene is sometimes referred to as the ‘phylogenetic likelihood’ [25], and is characterized by pmf Pr(**D**_*i*_|*G_i_*). Under the MSC, the probability of that gene tree given species tree *S* and population sizes ****N_e_**** is *f* (*G_i_*|*S, **N_e_***). Note that *S* = {*ϕ, **t**^n^*, **t**^s^}, where t^s^ is observed data and correspond to the sample ages (fossil ages and living taxa). Both *ϕ* and ***t**^n^* are parameters (random variables) and denote the species tree topology and internal node times, respectively.

The likelihood contribution to the species tree of a morphological character is captured by the phylogenetic likelihood Pr(**C**_*j*_|*S*) where **C**_*j*_ is the vector of states for the *j*-th character. The prior probability of the species tree under the FBD process is *f* (*S*|****θ****^FBD^), where ****θ****^FBD^ is a vector of FBD parameters. Finally, *f* (****θ****) describes the joint distribution over all parameters ****θ**** = {****θ****^FBD^*, **θ**^r^, **N_e_***} (where ***θ**^r^* denotes all remaining parameters not explicitly mentioned above). By combining the probability mass and density functions of all sampling distributions comprising the integrative model, we get the probability density of the species tree given the molecular, morphological and fossil age data:

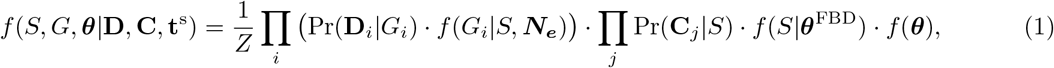

where *Z* = Pr(**D**, **C**) is the marginal likelihood, an unknown normalizing constant that does not need to be computed when using Markov chain Monte Carlo (MCMC) to sample from the posterior distribution.

When conducting inference under this model, MSAs are assumed to evolve along gene trees, which then inform the species tree via the MSC, whereas the morphological characters are assumed to have evolved along the species tree itself, and thus inform it directly. Ultimately both the MSAs and morphological characters inform the FBD parameters through the species tree (e.g., Supplementary Figs. S4, S5).

Under our integrative model, the likelihood of gene trees and of the discrete morphological tree (the latter being the species tree, *S*) are computed with a model in the generalized time reversible (GTR) family [26] and the Mk model [27], respectively. The MSC probability density is calculated based on species tree branch lengths, and on a function returning the effective population size for each branch. While in our analyses this function always returned a constant size within a branch, linearly changing population sizes are also supported by StarBEAST2 (and other functions like exponential or stepwise are also possible in principle). Lastly, the FBD prior assumes that the rates of speciation, extinction, and sampling of fossils are constant throughout the species tree.

#### Well-calibrated validation of model and operators

An integrative (hierarchical) Bayesian model like the one we introduce here consists of a collection of probability mass and density functions characterizing all likelihoods and priors. Although some components of this collection may have been individually validated, the collection itself needs to be validated as a whole. (Where new MCMC operators are introduced, they also need to be validated.) This type of validation can be seen as both an instantiation and probabilistic analog of what software engineers refer to as ‘integration testing’, whereby software modules are combined and tested together. This type of testing is a mandatory stage in the software development life cycle because it is often hard to predict how different modules will interact.

In the particular case of our FBD-MSC model, this is illustrated by the addition of fossils (via the FBD process) complicating the relationship between the species and gene trees under the MSC – when compared with a species tree model such as the simple birth-death process without fossils. The two key assumptions of a birth-death process that are relaxed under the FBD-MSC are (1) that the species tree must be ultrametric, i.e., each of the *n* species is sampled at a single point in time, and (2) that the number of nodes is fixed at (2*n* − 1) regardless of how the topology of the tree changes. The relaxation of both assumptions in the FBD-MSC required changes to the implementation of the MSC and related operators in StarBEAST2 (see the Supplementary Material for an example in Algorithm S3).

More specifically, relaxing the second assumption requires fundamental changes to the inferential algorithm. This is because previous implementations of MSC used Metropolis-Hastings MCMC, which does not allow for changing the number of dimensions in the model. But converting a fossil from terminal node to sampled ancestor will decrease (or in the other direction increase) the number of nodes and hence dimensions. In such cases, not only does an additional node age have to be estimated, but so do the parameters of the population size function of the corresponding branch. One possible strategy to sample the additional node age – the one we chose – is to use reversible-jump operators, such as those previously developed for BEAST 2 [12]. To sample the additional population size parameters, we implemented a composite model space approach [28–30] whereby population size parameters for the maximum number of species branches are being sampled, but only those corresponding to branches in the current topology are contributing to the likelihood.

Because our full model and related MCMC machinery are new in the ways described above, we verify correctness through a well-calibrated validation study. Here, many independent data sets are simulated under the full model (i.e., all pmf and pdfs), and correctness is deduced from appropriate posterior coverage upon MCMC chain convergence.

### Empirical analysis: the canid subfamily Caninae

The diverse family of dogs (Canidae) has a rich fossil record that has made this clade a model for ecological, evolutionary and methodological studies [31]. Canidae is comprised of three subfamilies – Borophaginae, Hesperocyoninae and Caninae – represented by carnivorans of jackal, fox and wolf semblance [32]. Borophaginae (∼66–69 species [32, 33]) and Hesperocyoninae (∼26–29 species [32, 34]) consist of only extinct species, with Caninae accounting for the remaining fossils (out of ∼140–178 [32, 35]) in addition to 36 living species [36].

Previous phylogenetic accounts of canids using morphology alone under the FBD model have shown that this type of data can produce sensible age estimates [37], but contrasting topologies, particularly in terms of root placement, when compared to molecular trees (in the case of Caninae [31, 38]). Here, we further examine the phylogeny of Caninae by carrying out an integrative statistical analysis where molecular and morphological data jointly inform the species tree.

### Compiling molecular data

To maximize the information available for phylogenetic reconstruction, we combined DNA sequences from five previous studies. Four of the studies contained segments of coding and/or non-coding DNA [39–42]. All sequences from the above studies were retrieved from GenBank (Supplementary Table S5). We excluded all sequences from loci other than nuclear autosomes. We also used only one segment for a given gene where multiple segments were available, avoiding segments from a study for which fewer taxa were available [42].

The fourth study included the coding sequences of multiple intron-less taste 2 receptor (*Tas2r*) genes [43]. After investigating these data, we identified and removed five sequences with likely erroneous labels, and three sequences that were probably either paralogs, degraded, or contained excessive errors. We also identified two pairs of sequences where the labels had likely been swapped, which we corrected (Supplementary Table S5). That investigation was partly based on a gene tree (Supplementary Fig. S12), inferred from the unaligned *Tas2r* sequences using `PASTA` [44] and available in Supplementary Material. Based on that gene tree we excluded the *Tas2R43* and *Tas2R44* genes, as four of the *Tas2R44* sequences appeared to actually be *Tas2R43* sequences. All *Tas2r* sequences were retrieved from GenBank, other than *Lycaon pictus* sequences, which were extracted from the supplementary material of that study (Supplementary Table S5).

The different data sets were somewhat heterogenous. For one study multiple representative specimens were sometimes available for one species [41], but not for other data sets. For another, multiple haplotypes were sometimes available for one specimen [39], but other data sets apparently used ambiguity codes to represent heterozygosity. To make the data sources more uniform, we chose one sequence per locus at random, and randomly resolved all ambiguity codes. At this stage we also excluded outgroup and domestic dog sequences.

For each locus, we aligned the corresponding sequences using `PRANK` [45]. The resulting MSA was trimmed using the “gappyout” method of `trimAl` [46]. Our final data set included 938 sequences from 58 loci and 31 extant Caninae species (out of the 36 known living species [36]). This means the amount of missing data, in terms of the number of sequences for a given taxon and locus that were not available, was 48% (Supplementary Fig. S13). A numerical summary of our molecular data set can be found in Supplementary Table S7.

### Compiling morphological and fossil data

Our morphological data set is derived from an existing character matrix [31]. Some of these characters were newly scored by the authors of that study but many had been published previously [33–35, 47–56]. This matrix included soft tissue, pigmental, ecological, developmental, behavioural, cytogenic and metabolic characters not available for fossil taxa. Since our interest in this data set is to use it for total-evidence inference, we retained 230 characters from the original matrix (indices 12 through 241 in the original study) corresponding to skull, dentition, body proportions and postcranial skeleton characters. These were generally available for both fossil and extant taxa.

Character states were available for all 31 extant taxa in our molecular data set, in addition to five extant or recently extinct species absent from our molecular data set (*Canis rufus*, *Dusicyon australis*, *Vulpes bengalensis*, *V. pallida* and *V. velox*) and 42 fossil taxa (out of ∼48–80 fossil species [32, 35]), for a total of 78 species. A total of 11,357 states were known, corresponding to a missing data rate of 37% in terms of the proportion of states which are unknown across all characters and species.

Stratigraphic ranges were extracted from the same publication as the character matrix [31], and the midpoint of each range used as the tip date of the corresponding taxa, rounded to the nearest 0.05 million years. If the range of a species reached the present, time zero was used as the tip date. While tip dates could also have been estimated, doing so would increase the number of parameters of our model, and potentially require longer MCMC chains; more importantly, without extensive expert curation it is not immediately clear which prior distributions to use for each of the different fossils in our data set – a problem similar to the characteristic node dating problem.

### Caninae analyses

We analyzed Caninae data under two models that treat molecular, morphological and temporal information as data; these are the aforementioned FBD-MSC and what we call the FBD-concatenation model. Both were identical in terms of most of the density functions characterizing their sampling distributions, the only difference being that for FBD-concatenation MSAs were assumed to evolve directly along the species tree (i.e., Π_*i*_ (Pr(**D**_*i*_|*G_i_*) · *f* (*G_i_*|*S, **N_e_***) is replaced with Pr(**D**|*S*) in Eq. 1, where **D** represents all MSAs after concatenation). Hence StarBEAST 2 operators were disabled for FBD-concatenation since gene trees are not part of that model; all other operators were shared. A full description of the model and prior choice used for the FBD-MSC analysis of Caninae is given in Supplementary Methods.

Four independent MCMC chains were run for each method, with 4,096 evenly spaced samples being collected. Individual chain lengths were 2^30^ (roughly one billion) states for FBD-MSC and 2^29^ (roughly half a billion) for FBD-concatenation. For each method, the chains were joined after discarding the first 64 samples (roughly 1.6%) from each chain as burn-in. Burn-in was determined by manual inspection of MCMC traces in Tracer 1.7 [57]. These chains were then thinned to one in every eight samples, for a total of 2,016 samples per method.

Summary statistics were calculated for each estimated distribution of trees using `DendroPy` [58]. These included the maximum clade credibility (MCC) tree, branch lengths, internal node ages and node support. For the purpose of calculating support values and ages, a node is defined as the root of a subtree containing all of, and only, a given set of extant taxa. Lineages-through-time (LTT) curves for FBD analyses were calculated using a custom script. Summary statistics and LTT plots were visualized using `ggplot2` [59] and `ggtree` [60].

## Results

### Correctness of the full model

We tested the correctness of our model implementation including the MCMC operators by repeating a validation procedure carried out in a previous study using the FBD model [10]. As in that study, we demonstrated the inference of correct tree topology probabilities by comparing our method to analytically derived probabilities (Table 1). We extended this validation by demonstrating the correct inference of internal node ages when compared to automatic integration using quadrature (Supplementary Fig. S11). Together these results indicate our MCMC operators have been correctly implemented and interact with the FBD model as expected.

**Table 1:** Number of replicate MCMC chains in which a particular topology was found within the 95% HPD interval (the expected number under a correct implementation is 95). Full details of operator configurations are provided in the Supplementary Material.

| Configuration \ Topology | T1 | T2 | T3 | T4 | T5 | T6 | T7 | T8 |
| --- | --- | --- | --- | --- | --- | --- | --- | --- |
| SA | 95 | 95 | 97 | 91 | 95 | 95 | 95 | 98 |
| UpDown | 94 | 97 | 95 | 97 | 96 | 95 | 94 | 93 |
| MSC | 92 | 96 | 95 | 97 | 93 | 95 | 95 | 93 |
| Coordinated | 96 | 95 | 93 | 95 | 90 | 95 | 97 | 97 |
| NodeReheight2 | 95 | 98 | 94 | 96 | 97 | 96 | 95 | 95 |
| Full | 91 | 92 | 94 | 96 | 97 | 98 | 99 | 96 |

To further confirm the correctness of our implementation, we carried out an extensive well-calibrated validation study (e.g., Zhang *et al*., Gaboriau *et al*. and Zhang, Drummond & Mendes [11, 61, 62]). This tests the implementation and illustrates statistical power and parameter identifiability. The simulations used for this study were conditioned on 40 extant taxa for each species tree, but covered a broad range of tree sizes when including fossil taxa (Supplementary Fig. S9).

As can be seen in Figure 2, all parameters had appropriate coverage, i.e., approximately 95% of 95%-HPD intervals contained the true simulated value. Furthermore, a high correlation between posterior estimates and true values was obtained for all parameters, with the exception of birth rate for which only a moderate correlation was observed. Apart from birth rates, the FBD-MSC had the power to estimate the other parameters in Figure 2 with useful precision. In the parameter space covered by our simulations, larger trees are likely necessary if one hopes to accurately estimate birth rates.

**Figure 2:**
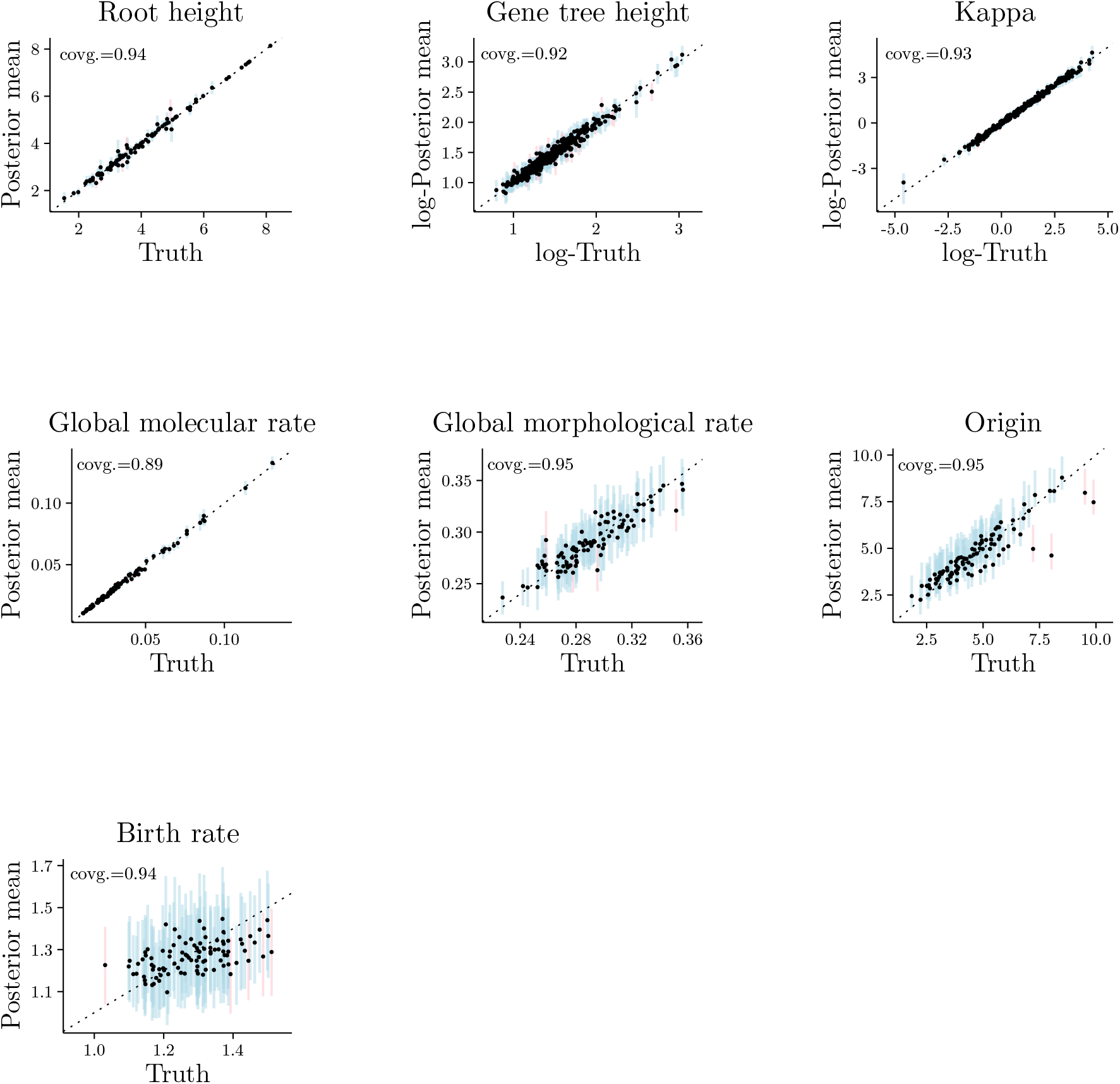
Parameter posterior means against their true simulated values, for 97 simulations (3 were excluded due to convergence issues). Blue lines correspond to the 95%-HPD intervals for a parameter, one line per simulation, when the true value was contained within the interval. When the true value was outside the interval, red lines were used instead. Panels for parameters “Kappa” and “Gene tree height” contain 388 data points (4 loci times 97 simulations). The coverage of each parameter is shown at the top-left corner of the corresponding panel.

Overall, results from our analytical validation and from Figure 2 allow us to conclude that our model (as well as MCMC proposal mechanisms and Hastings ratios) has been correctly implemented. Our conclusions are further strengthened by the fact that all the well-calibrated simulation code is independent of BEAST 2 code used in inference, and in large part written in different programming languages (Supplementary Table S4). Full details of the MCMC operators, methodology for the comparison of topology probabilities and internal node ages and methodology for the well-calibrated validation study are given in Supplementary Methods.

### Inferring time-trees of Caninae species

We summarized the posterior distributions of Caninae species trees inferred under FBD-concatenation and FBD-MSC models as maximum-clade-credibility (MCC) trees (Supplementary Figs. S14, S15). In order to compare branch length estimates from both models, we parsed their posterior distributions in two steps: (i) we pruned taxa providing only morphological data because their phylogenetic affinity is difficult to estimate, and (ii) we established a branch frequency threshold (see below) that branches had to meet in order to be compared. Both steps minimize topological differences in an attempt to make length comparisons fair; while arbitrary, these steps yielded posteriors whose trees had a reasonable number of branches inducing largely agreeing clades above and below.

A branch was compared only if both its parent and child nodes (the child clade is necessarily a subset of the parent clade) were present in at least 0.5% of the trees in both FBD-concatenation and FBD-MSC posteriors. Every terminal branch meeting this frequency threshold was estimated to be longer using FBD-concatenation compared with FBD-MSC, and internal branches present above the threshold in both distributions were generally inferred to be shorter (Fig. 3). In some cases the terminal branch lengths were inferred to be more than twice as long, e.g., those of *Urocyon* species.

**Figure 3:**
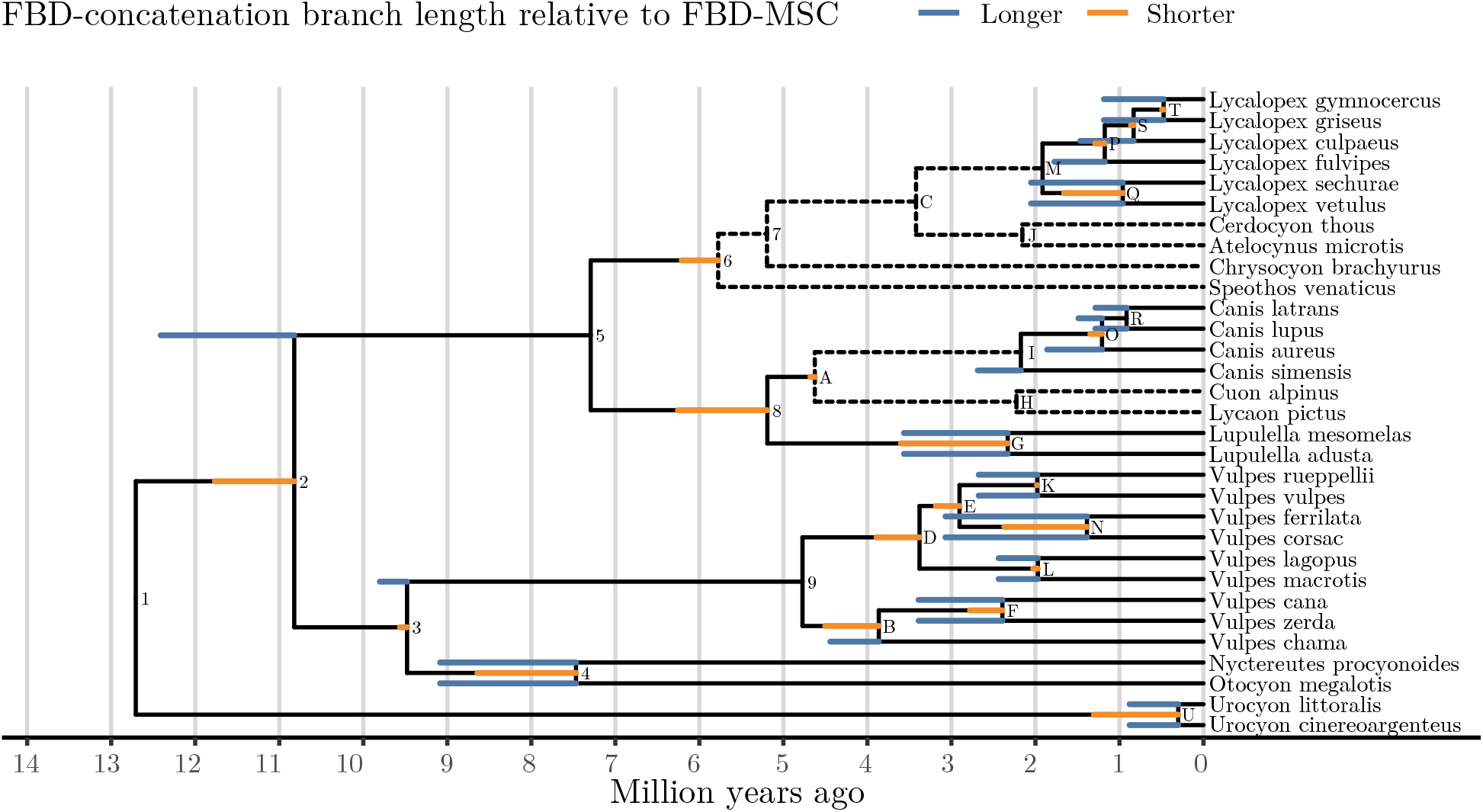
Branch length changes resulting from concatenation. The tree shown is the maximum clade credibility (MCC) tree with mean internal node ages from the FBD-MSC posterior distribution. When the length estimated by FBD-concatenation was longer than for FBD-MSC, the additional length is shown as an extension in blue. When the length was shorter, the reduction is shown as a truncation in orange. The difference in branch lengths is the mean among FBD-concatenation samples including that branch, less the FBD-MSC mean. Dashed lines represent branches not meeting the 0.5% frequency threshold described in the main text.

Several branches did not meet the frequency threshold, such as the branch connecting *Cuon alpinus*, *Lycaon pictus* and extant *Canis* up to their common ancestor. While *C. alpinus* and *L. pictus* belong to a clade sister to all extant and some stem *Canis* species in the full FBD-MSC MCC tree (Supplementary Fig. S14), *C. alpinus* is more closely related to extant *Canis* than it is to *L. pictus* in the full FBD-concatenation MCC tree (Supplementary Fig. S15). Under both models, the grouping of *Canis*, *Cuon* and *Lycaon* was well supported. The grouping of *Speothos* and *Chrysocyon* by FBD-concatenation was not well supported under the FBD-MSC, whose MCC tree shows *Speothos* as sister to the other extant South American canids (Supplementary Fig. S14, S15).

Mean effective population sizes (Table 2) were estimated well within the empirical range for canine species, which have highly variable census population sizes (Supplementary Table S8). We note that our model integrates out branch-specific *N_e_* values [63], and that what we estimated is the mean of an inverse gamma prior distribution on ****N_e_**** (see Supplementary Material for more details).

**Table 2:** Parameter estimates. All values are posterior mean estimates followed in brackets by the bounds of 95% highest-posterior-density intervals. ‘Molecular clock rate’ is the global (mean) rate that scales the relative locus-specific molecular rates. ^∗^expected number of character state changes (i.e., substitutions for molecular data) per million years. ^∗∗^the mean of the inverse gamma distribution fit to per-branch *N_e_g* values, which are effective population sizes *N_e_* scaled by generation time in millions of years *g* (see also Supplementary Table S8).

| Parameter | FBD-concatenation | FBD-MSc |
| --- | --- | --- |
| Molecular clock rate* ($\times 10^{-3}$) | 0.91 (0.74–1.11) | 0.62 (0.52–0.73) |
| Morphological clock rate* ($\times 10^{-2}$) | 3.11 (2.50–3.68) | 3.27 (2.69–3.78) |
| Mean population size** | NA | 1.51 (1.08–1.97) |
| Diversification rate ($\lambda - \mu$) | 0.09 (0.01–0.18) | 0.10 (0.01–0.19) |
| Turnover ($\mu \div \lambda$) | 0.85 (0.72–0.98) | 0.86 (0.73–0.99) |
| Sampling proportion ($\psi \div (\psi + \mu)$) | 0.16 (0.07–0.25) | 0.15 (0.07–0.25) |

### Concatenation estimates significantly larger molecular evolution rates

The mean posterior estimate of the overall molecular clock rate for (nuclear) protein-coding genes was 6.2 × 10^−4^ substitutions per site per million years using FBD-MSC, but a broader peak centered at the higher rate of 9.1 × 10^−4^ was observed using FBD-concatenation (Table 2, Fig. 4a). Our FBD-concatenation estimated average clock rate was consistent with previous rate estimates obtained without accounting for ILS, e.g., 10^−3^ for the RAG1 gene in mammals [64]. Unlike the molecular clock rates, however, the posterior distributions of the morphological clock rates largely overlapped between FBD-concatenation and FBD-MSC (Table 2, Fig. 4b).

**Figure 4:**
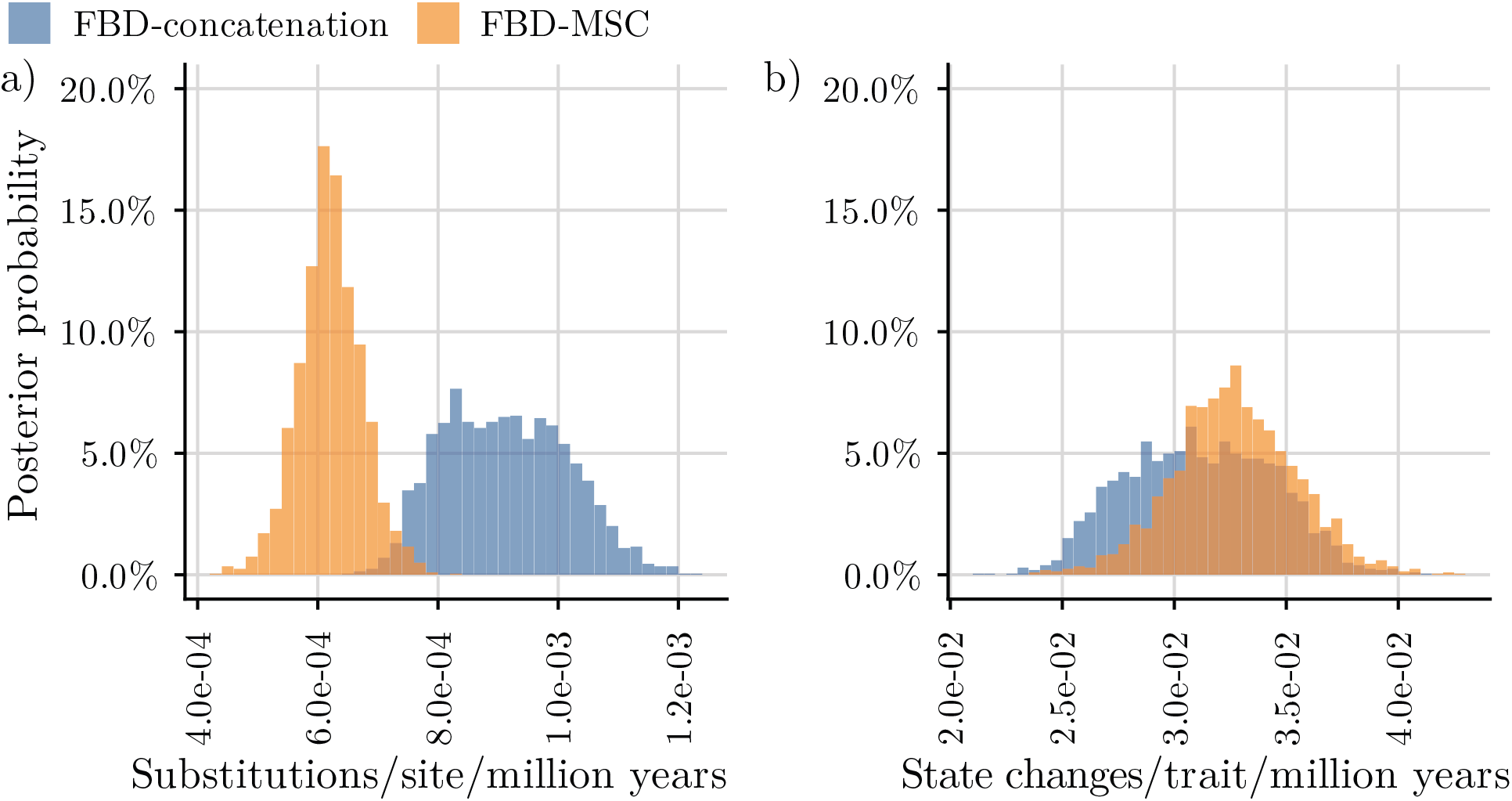
Posterior distribution of clock rates. Posterior probabilities of molecular clock rates (a) and morphological clock rates (b) were calculated using bin widths of 2 × 10^−5^ and 5 × 10^−4^ respectively.

Estimates of macroevolutionary parameters were also similar for FBD-MSC or FBD-concatenation (Table 2). We find the rate of extinction within Caninae to be high, with a lower bound on turnover of 72% using FBD-concatenation and 74% using FBD-MSC. This means the rate of extinction is at least 72% or 74% the rate of speciation for this subfamily.

### Caninae divergence time estimates are skewed under concatenation, but not under the MSC

For clades with crown ages younger than 4 million years ago in the FBD-MSC MCC tree, and with at least 0.5% posterior support using FBD-concatenation, the crown age was estimated to be older using FBD-concatenation, often with little overlap in the posterior distributions. The most extreme example is node N, the common ancestor of *Vulpes corsac* and *V. ferrilata*, inferred to be under 1.5 million years old using FBD-MSC, but around 3 million years old using FBD-concatenation (Fig. 5a).

**Figure 5:**
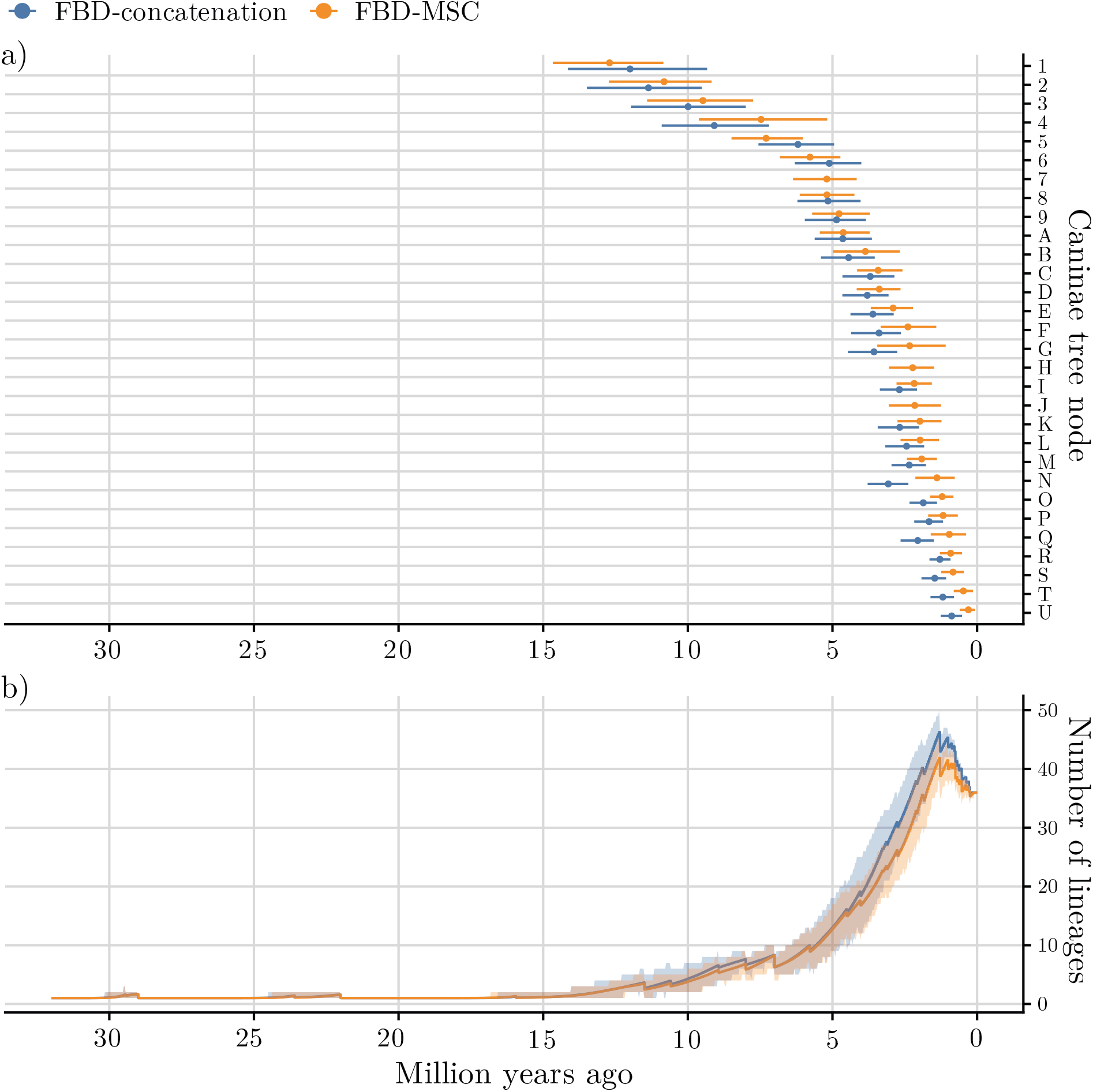
Tempo of Caninae evolution. Crown ages estimated by fossilized birth-death with multispecies coalescent (FBD-MSC) and with concatenation (FBD-concatenation) models (a), compared with lineages-through-time (LTT) curves including extinct lineages (b). Posterior mean internal node ages (solid circles) with 95% highest posterior density (HPD) intervals are estimated from samples where that clade is present after pruning all morphology-only taxa. Internal node labels correspond to those in Figure 3. Posterior mean estimates (solid lines) of LTT are calculated for 1,024 evenly spaced time steps spanning 0 to 32Ma. 95% highest posterior density (HPD) intervals calculated for each step are shown as ribbons.

Crown absolute ages estimated with FBD-concatenation for older clades were more in line with FBD-MSC estimates than those of younger clades; while posterior means could diverge, credible intervals were usually substantially overlapping (Fig. 5a). (When trees are measured in substitutions per site, however, deep nodes reappear as being older when estimated under FBD-concatenation, as a result of overestimated molecular clock rates; Supplementary Fig. S10.) Plotting estimated node log-ages under FBD-MSC as a function of those from FBD-concatenation (Fig. 6) revealed that younger nodes were consistently estimated as older by concatenation, so much so that speciation events within the past 500 thousand years were not inferred with this approach.

**Figure 6:**
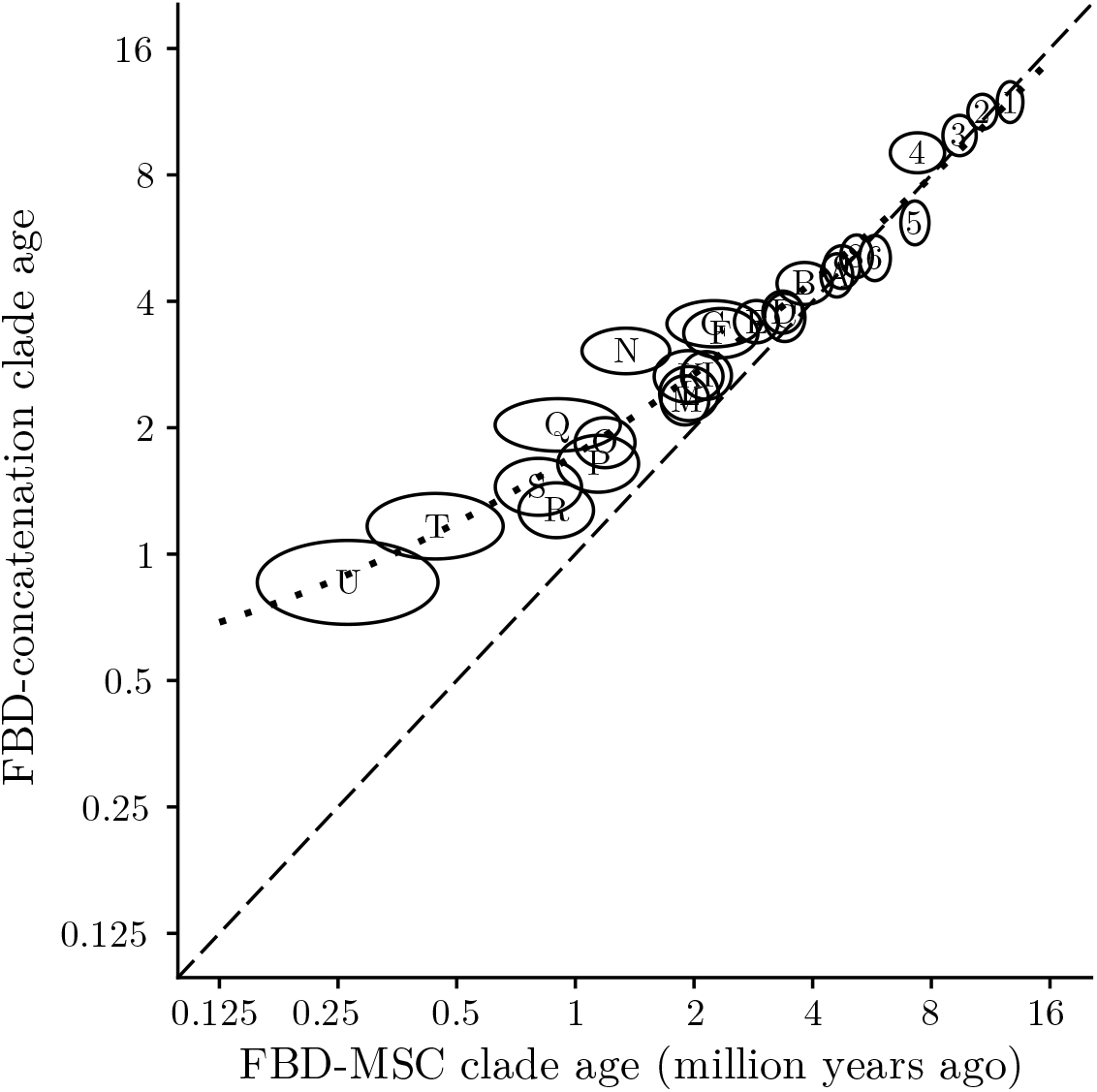
Correlation between log-node ages from the posterior distributions of species trees pruned of morphology-only taxa. Internal nodes from the pruned FBD-MSC MCC tree are drawn as ellipses with their labels from Fig. 3. Ellipses are centered on the mean estimate of the log-node age for both methods. The width and height of each ellipse corresponds to the standard deviation of the log-node ages for FBD-MSC and FBD-concatenation respectively. The dashed black line shows the 1:1 line along which estimates are equal, and the dotted line is the quadratic line of best fit.

The skewed ages of younger nodes inferred using FBD-concatenation will affect macroevolutionary analyses, including analyses of lineages-through-time (LTT). To demonstrate this, we computed LTT curves based on the species tree posterior distributions inferred using FBD-concatenation and FBD-MSC (Fig. 5b). For both methods the curves are concave upwards, as expected for a birth-death model of evolution with good taxon sampling [65]. However the curves diverge towards the present, so that the burst of speciation leading to the abundance of extant Caninae species is estimated to occur earlier using FBD-concatenation compared with FBD-MSC.

### Computational performance

We report performance summaries for all Caninae analyses in Table 3; these were all run on an 4GHz Intel i7-8086K CPU. The lowest effective sample sizes were observed for the coalescent probability density and the phylogenetic likelihood of the *T2R42* locus under the FBD-MSC and FBD-concatenation, respectively. From our observations, tip-dating methods will require substantially longer chain lengths than node-dating methods, but using the MSC to model gene tree evolution does not incur a substantial computational performance penalty. For each method four chains were run in parallel, so the actual time spent waiting on chains to finish was roughly one quarter the combined CPU hours.

**Table 3:** Computational performance of Caninae analyses under the two models.

| Model | Combined CPU hours | Min. ESS | ESS per hour |
| --- | --- | --- | --- |
| FBD-MSc | 399 | 274 | 0.69 |
| FBD-concatenation | 326 | 229 | 0.70 |

A previous study on MSC inference using MCMC found that doubling the number of loci increases the time to 200 ESS roughly 7-fold [20]. Thus even by running multiple chains in parallel as done in this study, we anticipate our MCMC-based implementation will be applied to data sets of fewer than 1,000 loci.

## Discussion

We introduced a new integrative model, the FBD-MSC, for total-evidence analysis of data from extinct and extant species. Because we model the population-level phenomenon of ILS under the FBD-MSC, we make it possible for comparative biologists to carry out statistical inference across evolutionary time scales for the first time.

We also carried out critical validation of the FBD-MSC model and related operators, in a thorough validation study. While in an ideal scenario there should be no surprises when previously validated models are combined, there are often fundamental conceptual consequences for combining sampling distributions that only become apparent once we embark on building the composite model.

All previous descriptions of the MSC (e.g., Heled & Drummond, Pamilo & Nei and Rannala & Yang [63, 66, 67]), for example, have assumed the species tree is ultrametric. By combining the MSC with the FBD, a number of assumptions that were valid for ultrametric species trees no longer held. The most glaring idiosyncrasy of the FBD-MSC is that the number of branches, and therefore the number of population sizes, becomes a random variable of the composite model. As outlined in the above sections, there are multiple technical solutions to this problem. Further extensions of our integrative model could introduce an oriented species tree formalism [68], which could allow the population size to remain the same through successive fossils from the same morphological species.

These considerations highlight the point that the construction of novel joint models often requires, and leads to, new thinking. Their implementation needs more than just good book-keeping. As the size and complexity of phylogenetic models continues to grow, so too should we expect to discover emergent properties. We believe that careful construction and validation of joint models is a fundamental contributor to scientific novelty in molecular evolution and phylogenetics.

### Caninae taxonomy

We tested our new method – and compared it to the popular alternative of concatenation – with a data set of molecular and morphological characters of canine fossil and living taxa. We still do not fully understand how concatenation (of both molecular and morphological characters) can bias species tree inference by not capturing the possible topological independence between characters. Unlike the MSC, concatenation “channels” the information from all characters into supporting a single tree topology, which is then taken as a proxy for the species tree. Under the MSC, conversely, sets of sites are allowed to evolve along their own gene trees, whose topology might on average be less resolved than the single tree proxy estimated through concatenation. These more uncertain gene trees must then inform the species tree through the MSC density function. Morphological characters can thus perhaps be seen as having relatively greater influence in FBD-MSC inference when compared to FBD-concatenation.

Indeed, support for the topology we obtained with the FBD-MSC is echoed by morphological phylogenetic studies of Caninae [52, 54], probably as a result of their specialized dentitions. A previous study of Canidae (the family to which Caninae belongs) that combined morphological characters and mitochondrial sequence alignments found that support for (*L. pictus*, *C. alpinus*), for example, came only from the morphological data, and proposed that the responsible characters are likely convergent due to the hypercarnivory of these two species [38]. It could be due to the lower relative signal of morphological characters (with respect to nucleotide sites) under FBD-concatenation that the aforementioned clade is not recovered. Similarly, the inference of *Speothos* as sister to the other extant South American canids under the FBD-MSC model might reflect the same phenomenon. These results suggest that improved morphological models will be a fruitful avenue of ongoing research.

### Caninae evolutionary rates and divergence times

When analyzing our data sets with concatenation, we observed this method estimated markedly larger global (mean) molecular evolutionary rates than the FBD-MSC. We expect this outcome for at least two reasons. First, concatenation treats coalescent times as speciation times, when the former must always be at least equal, but usually greater than the latter. As a result, estimated tree lengths (and internal node ages) in substitutions per site under FBD-concatenation will be larger than under FBD-MSC. The reconciliation of the calendar ages of deeper nodes with the fossil record then manifests as a higher overall molecular rate, a phenomenon that in the context of morphological models was dubbed the “deep coalescence effect” [69]. Second, molecular rates are spuriously inflated as a result of sites within the concatenated alignment having different genealogies because of ILS, which leads to hemiplasy [17].

Concatenation also estimated larger terminal branch lengths (i.e., older divergence times of contemporary species). Again, we believe this is due to internal nodes under this procedure representing coalescent times rather than speciation times. Under the MSC, coalescent times are always deeper than corresponding speciation times. Furthermore, terminal branch lengths can be inferred to be larger because of hemiplasy when concatenation is carried out in the presence of ILS [17]. Irrespective of the cause, these results suggest that using the FBD-MSC model can significantly improve branch estimation from real data.

Our findings agree with a previous empirical study that also demonstrated concatenation can lead to longer terminal branch lengths relative to the MSC model [70]. That study was nonetheless limited to molecular data from extant species and was therefore missing critical dating information that only serially timed data (like fossils) can provide, and that can only be incorporated through an integrative model like ours. We note that while it is possible that FBD-concatenation is correctly estimating the empirical molecular rate (the truth is unknown), with FBD-MSC underestimating it, we find this unlikely. Previous theoretical and simulation work reveal biases in line with what we observed here, and that these biases are corrected (or should be corrected) by the MSC as a result of modeling population-level processes [17, 21].

Finally, another novel finding included the fact that estimates of younger node divergence times could be substantially biased upwards by concatenation, but those of deeper nodes were less affected. The causes for this result are likely manifold. Mendes & Hahn [17] showed that hemiplasy spuriously increases terminal branch rates while decreasing internal branch rates, possibly to a lesser degree when ILS happens among more species (see Supplementary Fig. 5 from that study). In the absence of calibrations, this effect translates to longer terminal branches and shorter internal branches. Because a larger number of (shortened) internal branches is expected between deeper nodes and the present, the net effect of ILS on deeper node ages should be less pronounced. Future studies may examine the contribution of ILS to the convergence of FBD-concatenation and FBD-MSC estimates at deeper time scales, and its interplay with tip dating and different molecular clock models.

## Conclusion

We have demonstrated that failing to model population-level processes when inferring species trees using an FBD model will substantially shift estimates of branch lengths, species divergence times and clock rates, and exclude the possibility of recent speciation. This is the first time that such biases are quantified in a real data set, as well as addressed using superior modeling. Our newly implemented FBD-MSC model accounts for the coalescent process while still being powerful enough to precisely recover rates and times. We also found topological differences between FBD-MSC and FBD-concatenation, but these may be due to traits being treated independently, when they can evolve in concert, e.g., towards hypercarnivory. New models could either rule in or out the *Lycaon*+*Cuon* grouping by ascribing their similar morphology to homology or convergent evolution. Alternatively, support for this putative clade could be further scrutinized through expanded sampling of fossil taxa, traits or genes.

A number of other avenues for further development of FBD-MSC are open. These include accounting for gene flow after speciation [71] or between lineages [72, 73]. Recent advances in speciation models beyond cladogenesis can also be applied within an FBD-MSC framework, for example treating species as a kind of trait evolving along a population tree [74], or incorporating budding speciation [68]. The modular architecture of BEAST 2 makes FBD-MSC analyses with the many substitution models in BEAST 2 immediately available, and will make future extensions as in the examples above relatively straightforward to implement. As it is, our method can be used today to infer time-scaled species trees and rates through total-evidence tip dating, without the problems caused by concatenation.

## Acknowledgements

We thank Craig Moritz for his advice on preparing the manuscript, and the late Colin Groves for his insight into the Caninae fossil record. Graham Slater provided expertise and insight regarding Caninae taxonomy. Constructive comments from Jeet Sukumaran, David Swofford and Gavin Naylor helped us improve the rigor and clarity of our manuscript. This work was supported by the Royal Society of New Zealand (16-UOA-277 to FKM, AJD, DW, NJM, TGV and TS); the Australian Research Council (FL110100104 to Craig Moritz supported HAO); and the European Research Council (PhyPD:335529 to TS).

## Notes

### Competing Interest Statement

The authors have declared no competing interest.

### Summary of Updates

The text has undergone substantial revisions for clarity, and we have added an analysis of node dating using the multispecies coalescent to the supplementary materials.

