## Supplementary Material for "Novel Integrative Modeling of Molecules and Morphology across Evolutionary Timescales"

### Supplementary information for “Novel Integrative Modeling of Molecules and Morphology across Evolutionary Timescales”

[†]These authors contributed equally.

### Supplementary methods

#### Previously developed MCMC operators

Apart from `UpDown`, which is used to change both species and gene trees, BEAST 2 tree operators are only used to change the gene trees in our FBD-MSC implementation. These operators are `TreeScaler`, `TreeRootScaler`, `Uniform`, `SubtreeSlide`, `Narrow`, `Wide`, and `WilsonBalding`. The BEAST 2 `Scale` operator is used to change the other real parameters of the model, e.g., the origin height. These operators have been previously described [1].

Sampled Ancestor operators are used to change a species tree potentially containing sampled ancestor nodes, for which in-degree = out-degree = 1. They are `LeafToSampledAncestorJump`, `SATreeScaler`, `SATreeRootScaler`, `SAUniform`, `SANarrow`, `SAWide` and `SAWilsonBalding`. These operators have been previously described [2].

StarBEAST2 operators make coordinated changes to the heights of species and gene tree nodes. `CoordinatedUniform` changes the height of a single non-root internal species tree node, and `CoordinatedExponential` changes the height of the species tree root node. Both operators have been previously described [3]. These operators have been modified to only change the height of nodes corresponding to true bifurcations in the species tree, i.e., not a sampled-ancestor node.

#### The `TREE SLIDE` operator

In addition to `CoordinatedUniform` and `CoordinatedExponential`, StarBEAST2 improves on the `TREE SLIDE` [4] operator and renames it `NodeReheight2` after a similar operator that only works on ultrametric trees. The renaming was done for backward compatibility with previous versions of StarBEAST2. This improvement took place with StarBEAST2 v15.5, in which `TREE SLIDE` was reimplemented from scratch and made compatible with fossil/ancient taxa including sampled ancestors. Below we describe the development of this operator.

When a rooted oriented tree is traversed preorder, postorder, or (for binary trees) inorder, this will generate a linear sequence of nodes in a deterministic order. In this document we will let $s = f(\mathbb{T})$ denote the linear sequence of nodes derived from inorder traversal of any rooted oriented binary tree $\mathbb{T}$. It was previously demonstrated that an ultrametric oriented binary tree $\mathbb{T}$ can be reconstructed from the sequence of leaf labels $\sigma$ which are in the same relative order as their corresponding nodes in $s$, together with the vector $a$ of most recent common ancestor (MRCA) heights for each pair of leaves which are immediately adjacent within $\sigma$ [5].

For example, the top-left tree in Fig. S1 can be transformed into $\sigma = [A, B, C, D], a = [9, 6, 12]$. Notice that leaves and MRCA nodes are interleaved in $s$, so that each immediately adjacent pair of nodes in $\sigma$ are the $i^{th}$ and $i^{th} + 2$ nodes visited for each odd value of $i$, and the MRCA node for that pair is the $i^{th} + 1$ node visited. All intervals and indices used in this document will be closed and 1-based. The tree can be reconstructed following a previously developed algorithm [6], an example of which is shown in Fig. S1.

Because trees with the same topology and times but different orientations are nonidentifiable using many popular phylogenetic models (e.g., general time reversible substitution models), many phylogenetic methods including StarBEAST2 treat trees as unoriented. However unoriented trees can still be transformed and reconstructed by randomly choosing a left–right orientation of children at each internal node.

Based on the relationship $\mathbb{T} \mapsto (\sigma, a)$, a symmetric MCMC proposal for an *unoriented* binary ultrametric tree $T$ was

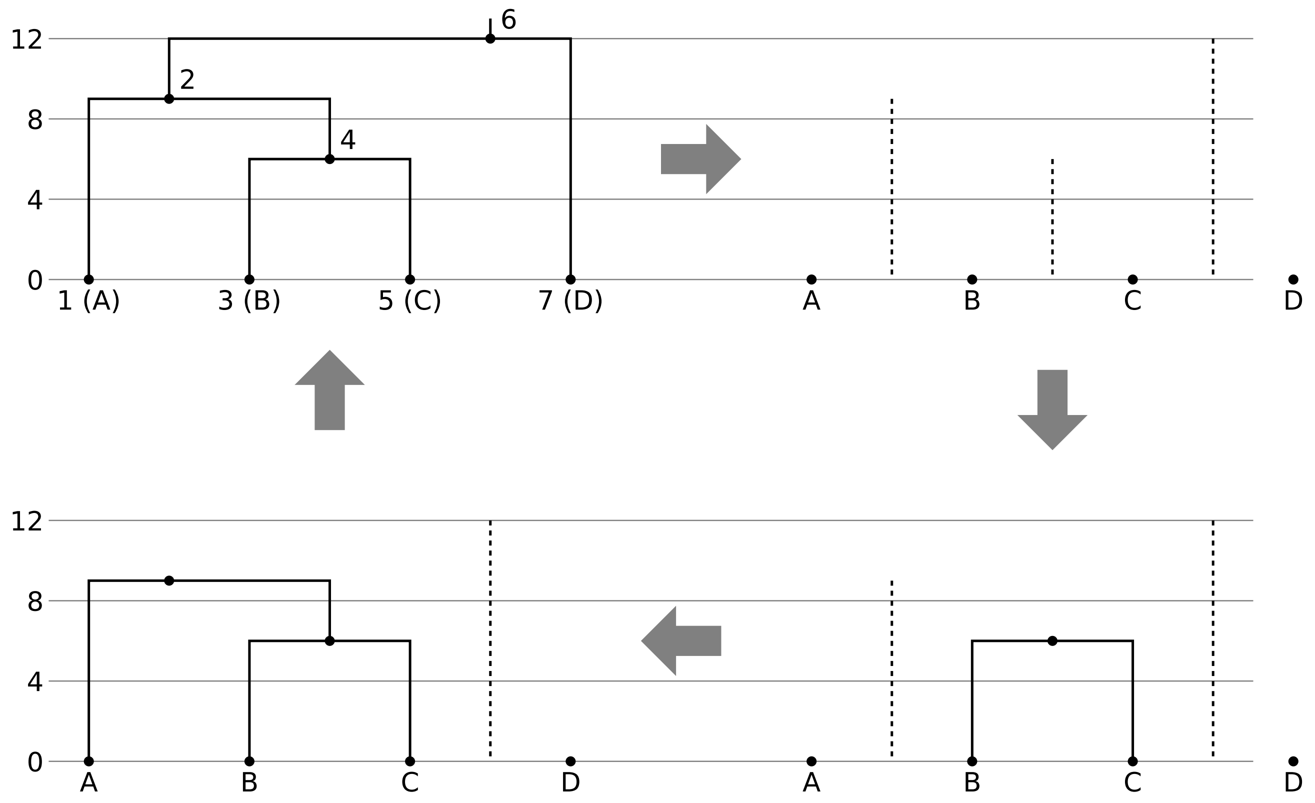


Figure S1: Transformation and reconstruction of an ultrametric oriented tree. Each node in the tree $\mathbb{T}$ is numbered according to its index in $s$, and leaf nodes labeled A through D. The tree is transformed to the leaf order $\sigma$ and MRCA heights $a$ (top-left to top-right). An internal node is created for each height in order of lowest to highest, the horizontally closest orphan (parentless node) to its left is made the left child, and the closest orphan to its right is made the right child (top-right clockwise to top-left).

introduced [5]. First, a random left–right orientation of children is chosen for each internal node to determine the order of $\sigma$ and $a$. Second, a new vector of MRCA heights $a^*$ is generated from $a$ using a random walk with reflection to avoid negative values. The proposed tree $T^*$, which will have different heights and may also be different in topology from $T$ due to the new heights, is reconstructed from $(\sigma, a^*)$. An example implementation is Algorithm S1.

The proposal is symmetric for two reasons. One, because the orientation of the tree is sampled uniformly at random from one of the $2^{n-1}$ possible orientations where $n$ is the number of taxa. Two, because reflection of a uniform probability mass is used to choose each new MRCA, the perturbation of node heights is symmetric [5].

`TREE SLIDE` is based on the MCMC proposal described above, but uses recursion to reconstruct the tree, and no longer uses a tunable parameter or random walk to modify $a$. Instead it chooses a new height between 0 and a natural upper bound uniformly at random for a single randomly picked internal node. When the multispecies coalescent (MSC) model is used and assuming that each species has at least one sampled allele, a natural upper bound for the height of a given node can be derived from the coalescent times.

The internal nodes of gene trees represent coalescent events, and the leaves represent sampled alleles. For an internal gene tree node $x$, the set of sampled alleles represented by the leaves located in the subtree defined by the left child branch of $x$ is denoted $x_l$. Likewise the set of sampled alleles represented by leaves descended from the right child branch is denoted $x_r$. Which child branch is left and which is right is arbitrary, so we can typically use the left–right orientation

**Algorithm S1:** Example implementation of the original MCMC proposal.

```
Data: current unoriented ultrametric binary tree T of n taxa
Result: proposed unoriented ultrametric binary tree T* of n taxa
Tunable parameter: δ
Function Inorder-Traversal(node v, linked lists σ and a) is
    if v is an internal node then
        flip a coin to choose the left–right order of children of v;
        Inorder-Traversal(left child of v, σ, a);
        append the height of v to a;
        Inorder-Traversal(right child of v, σ, a);
    else
        append the label of v to σ;
    end
end
let σ, a, a* and s be empty linked lists;
Inorder-Traversal(the root node of T, σ, a);
for i = 1 to n − 1 do
    let h ~ uniform(a_i − δ, a_i + δ);
    append |h| to a*;
end
for i = 1 to n do
    append a new leaf node with the label σ_i and height 0 to s;
    if i < n then
        append a new internal node of height a*_i to s;
    end
end
for i = 1 to n − 1 do
    let j be the index of the i^th lowest internal node in s;
    let l be the orphan with the index in s nearest to but less than j;
    let r be the orphan with the index in s nearest to but greater than j;
    set l as the left child of s_j;
    set r as the right child of s_j;
end
initialize and return T* using the highest internal node in s as the root;
```

implicit in whatever data structure is used to store the tree.

When the height of an internal node $X$ from an oriented species tree $\mathbb{T}$ is changed, the new height must be below any coalescence between alleles sampled from species to the left in of $X$ in $s = f(\mathbb{T})$ with species to the right. There is a mapping of sampled alleles to species, so we can define the sets of sampled alleles $X_l$ and $X_r$ which map to the sets $X_L$ and $X_R$ which are the species to the left and to the right respectively. Any gene tree internal node representing a coalescence between an $X_L$ and an $X_R$ species will meet both these conditions:

1. $x_l \cap X_l \neq \varnothing$

2. $x_r \cap X_r \neq \varnothing$

Or if not meeting the above, because the orientation of the gene trees should not change the result, will meet both these conditions:

1. $x_r \cap X_l \neq \varnothing$

2. $x_l \cap X_r \neq \varnothing$

A natural upper bound of the height of $X$ is therefore the smallest value among the heights of gene tree nodes meeting the first or second set of conditions above, as all MRCA heights will remain below the corresponding coalescent events. The proposed species tree will therefore remain compatible with all the gene trees, and unlike naïve tree proposals, `TREE SLIDE` will never be rejected for violating MSC constraints.

`TREE SLIDE` restricts the new node height to between the bounds described above (an example implementation is shown in Algorithm S2). Because the node height is chosen uniformly at random, it remains a symmetric move. The recursive algorithm to reconstruct the species tree was described previously [4], and an example proposal is shown in Fig. S2. In the example proposal, $X_L = [A]$, $X_R = [B, C, D]$, $X_l = [a]$ and $X_r = [b, c, d]$. For the lower constraining gene tree node, $x_l = [a]$ and $x_r = [b]$. For the higher constraining gene tree node, $x_l = [a, b]$ and $x_r = [c, d]$. For the non-constraining gene tree node, $x_l = [c]$ and $x_r = [d]$.

The `TREE SLIDE` `Reconstruct-Tree` function (Algorithm S2) recursively segments $s$ into left and right intervals. Each interval will correspond to a subtree, and the function identifies the highest node within an interval as the root of that subtree. The recursion begins with an interval spanning the whole of $s$, which corresponds to the entire tree $\mathbb{T}$ and the highest node is in fact the root of $\mathbb{T}$. Branches are drawn from the root of a child subtree to the root of its parent subtree (Fig. S2).

This operator was renamed `NodeReheight` in BEAST 2 [7], and renamed `NodeReheight2` after being optimized for computational performance in StarBEAST2 [3].

The fusion of the FBD and MSC models means that `TREE SLIDE` type proposals may generate invalid species trees, or may not be able to function at all. For our implementation of the FBD–MSC in StarBEAST2, we modified `TREE SLIDE` to address all issues. Each issue, and how we addressed it, is described below.

##### Serial sampling

Serially sampled trees can no longer be reconstructed from their inorder sequence of leaf labels $\sigma$ and MRCA heights $a$ alone, because the heights of leaf nodes which are older than the present are not stored by that representation. For serially sampled trees, we need an additional vector of heights $b$ to store the heights of leaf nodes in the same order those leaves appear in the inorder traversal sequence $s$.

Also because of serial sampling, the lower bound for a given internal node height may be greater than zero. Any node will have some number of descendent nodes (leaves and possibly other internal nodes) through its left child branch and some other number through its right child branch, which we will call $y$ and $z$ respectively. For an internal node in a binary tree, $y \geq 1$ and $z \geq 1$. For the internal node $X = s_i$ in $s = f(T)$, its left descendents will be $X_L = s_{i-y,\dots,i-1}$ and its right descendents will be $X_R = s_{i+1,\dots,i+z}$.

Since $s$ must be an ancestor of $s_{i-1}$ and $s_{i+1}$, its height must be greater than either of those two descendents. These are always leaf nodes because of the interleaving of leaf and internal nodes in $s$, but under the FBD–MSC model any leaf may have a greater than zero node height. Therefore to avoid proposing invalid trees, our new operator uses the larger height of $s_{i-1}$ and $s_{i+1}$ as the lower bound for the new height of $X$.

Extensive testing has revealed that ignoring the height of nodes other than $s_{i-1}$ and $s_{i+1}$ does not result in an invalid

**Algorithm S2:** Example implementation of `TREE SLIDE`, with changes from Algorithm S1 highlighted in red.

```
Data: current unoriented ultrametric binary tree T of n taxa
Data: embedded gene trees G
Result: proposed unoriented ultrametric binary tree T* of n taxa
Function Inorder-Traversal(node v, linked lists σ and a) is
    if v is an internal node then
        flip a coin to choose the left–right order of children of v;
        Inorder-Traversal(left child of v, σ, a);
        append the height of v to a;
        Inorder-Traversal(right child of v, σ, a);
    else
        append the label of v to σ;
    end
end
Function Reconstruct-Tree(linked list of nodes s of length m) is
    let i be the index of the highest node in s;
    if i is the index of an internal node then
        set the left child of s_i to Reconstruct-Tree(s_{1,...,i−1});
        set the right child of s_i to Reconstruct-Tree(s_{i+1,...,m});
    end
    return s_i;
end
let σ, a, a* and s be empty linked lists;
Inorder-Traversal(the root node of T, σ, a);
/* j points to the MRCA height to change and is also the length of X_L */
let j be a random integer between 1 and n − 1;
let X_l be the sampled alleles which map to X_L = σ_{1,...,j};
let X_r be the sampled alleles which map to X_R = σ_{j+1,...,n};
let upperBound = ∞;
for x ∈ g ∈ G do
    let x_l be the sampled alleles descended from the left child branch of x;
    let x_r be the sampled alleles descended from the right child branch of x;
    if (x_r ∩ X_r ≠ ∅ and x_l ∩ X_l ≠ ∅) or (x_r ∩ X_l ≠ ∅ and x_l ∩ X_r ≠ ∅) then
        set upperBound to the height of x if this lowers upperBound
    end
end
foreach e ∈ a do append e to a*;
let h ∼ uniform(0, upperBound);
change a*_j to equal h;
for i = 1 to n do
    append a new leaf node with the label σ_i and height 0 to s;
    if i < n then
        append a new internal node of height a*_i to s;
    end
end
initialize and return T* using Reconstruct-Tree(s) as the root;
```

tree being proposed.

##### Infinite upper bound

Under the FBD–MSC model, there may be no constraining coalescent events for a given node if there is only morphological data and no molecular loci for some species. This will be the case if for a species tree node $X$ in $s$, there are no molecular

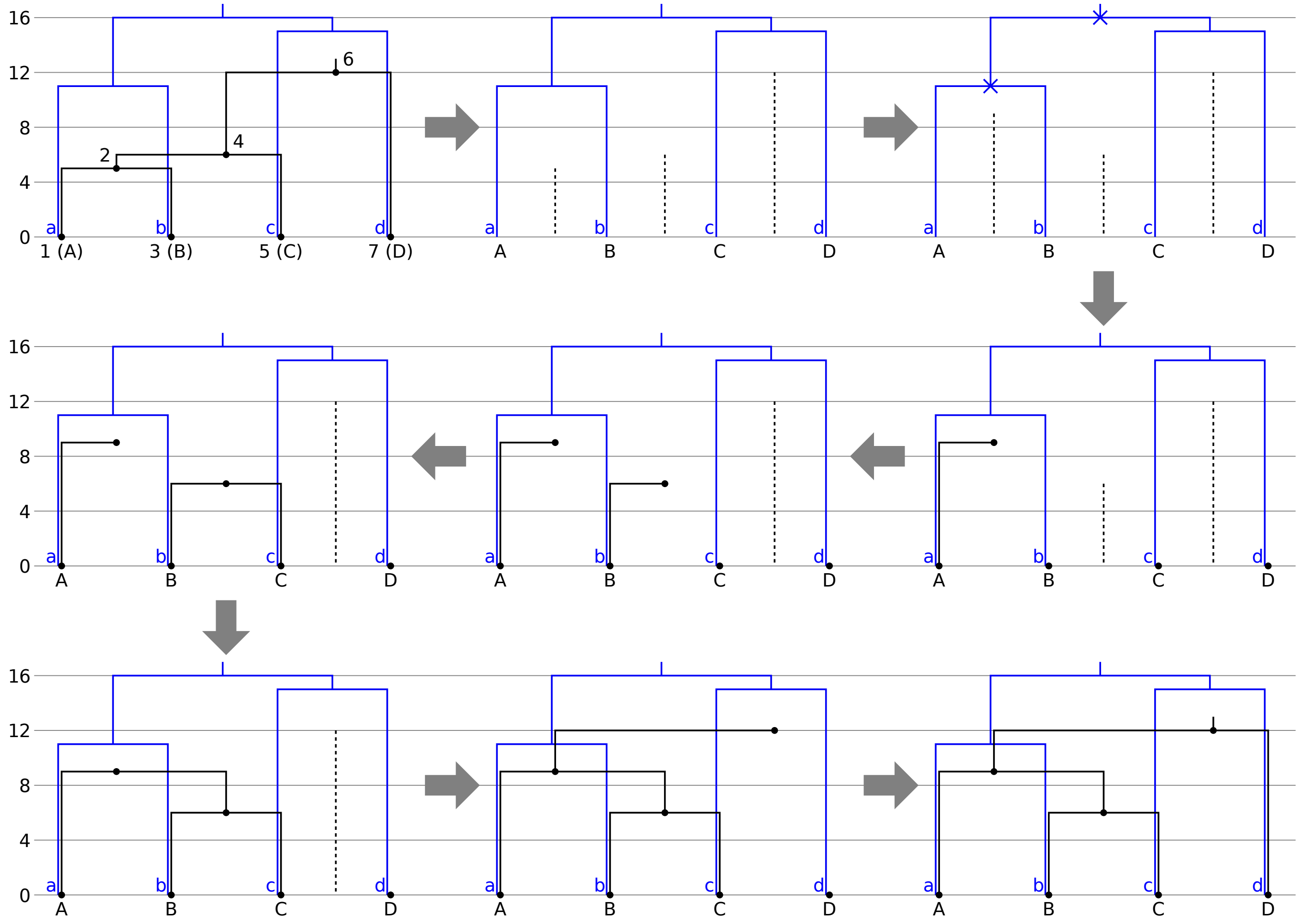


Figure S2: Example TREE SLIDE proposal using a single gene tree. Extant species are labeled A through D, and their corresponding sampled alleles are labeled a through d. The species tree is transformed to the leaf order $\sigma$ and MRCA heights $a$ (top-left to top-right). One height is modified to between 0 and the lowest height among coalescent events (blue crosses) linking alleles on the left with those on the right of the new height (top-right to middle-right). The species tree is then reconstructed recursively as described in Algorithm S2 (counter-clockwise middle-right to bottom-right).

data for $X_L$, and/or if there are no molecular data for $X_R$. In other words, if $X_l = \varnothing$ or $X_r = \varnothing$.

The FBD origin can be used as a natural upper bound, but if the FBD process is conditioned on the root instead of origin height, there will be no explicitly sampled origin to condition on. In order to make TREE SLIDE function without a natural finite upper bound, we still pick an internal bifurcation at random to change [4], but go back to using a random walk with reflection [5].

The upper bound for $X$ will be the lowest coalescence between $X_l$ and $X_r$ if this is available, the origin height if it is available, or whichever is smaller if both are available. If neither is available the upper bound will be infinity, as it is for the original MCMC operator [5]. To arrive at the proposed node height, the change in node height $\delta$ will be reflected against the lower and upper bounds until the proposed node height is within bounds.

#### Sampled ancestors

Under the FBD model, fossil taxa may be sampled ancestors with one parent and one child. However the current FBD implementation in BEAST assumes a binary tree data structure, where nodes are either internal with two children or leaves with none. To accomodate sampled ancestors, they are stored as a combination of two nodes; a leaf representing the sampled ancestor, and a "fake" internal node. The sampled ancestor is a child of the fake node via a zero length branch. The grandparent and sibling of the sampled ancestor in the binary tree are the actual parent and child respectively of the sampled ancestor under the FBD model.

This binary tree representation conveniently makes it possible, with a few tweaks, for `TREE SLIDE` to work with sampled ancestors. First of all, this operator is not intended to sample the heights of fossil taxa (including sampled ancestors). So when a height in $a$ is picked to be changed, this choice is restricted to elements of $a$ which correspond to true bifurcations (not leaves or fake nodes).

Second, the recursive algorithm in `TREE SLIDE` to reconstruct the species tree uses the highest node in a segment of $s$ as the root of a subtree. Since sampled-ancestor leaf nodes are the same height as their fake node parent nodes, to avoid choosing a sampled ancestor leaf when reconstructing the tree, this algorithm was modified to always choose an internal node (a true bifurcation or a fake node) as the subtree root, except when there is only one node in the current segment of $s$ (Algorithm S3).

Third, when the midpoint of stratigraphic ranges are used to date fossils, multiple fossils may be *a priori* assigned the same time-before-present and their nodes will occupy the same height in the tree. When two fossils are both sampled ancestors with the same height along the same lineage, we call them "superimposed" because they exist in exactly the same point in the tree under the FBD model. Other species tree operators used by StarBEAST2 are incompatible with superimposed nodes, but we have not yet found a way to determine in advance whether changing a given node height by a particular amount will lead to superimposed nodes (Fig. S3). So instead we check whether nodes are superimposed as the tree is reconstructed, and if they are, pre-emptively reject the move before calculating the acceptance ratio.

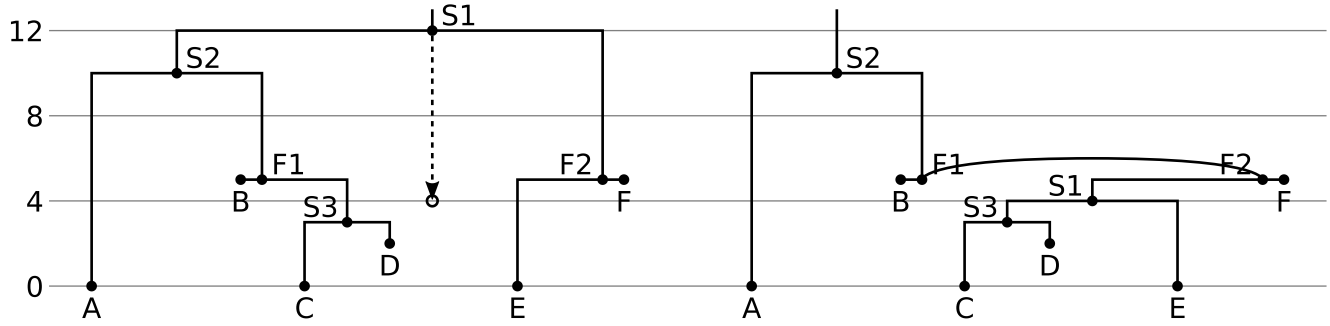


Figure S3: Changing an MRCA height may lead to superimposed nodes. Leaves and sampled ancestors are labeled A through F. True bifurcations are labeled S1 through S3. fake nodes are labeled F1 and F2. By moving the MRCA height corresponding to the S1 node to below the height of F1 and F2 (left), the fake nodes will be superimposed in the reconstructed tree (right).

**Algorithm S3:** FBD compatible implementation, with changes from Algorithm S2 highlighted in red.

```
Data: current unoriented serially sampled tree T of n taxa
Data: optional origin O
Data: embedded gene trees G
Tunable parameter: δ
Result: proposed unoriented serially sampled tree T* of n taxa
Function Inorder-Traversal(node v, linked lists σ, a and b) is
    if v is an internal node then
        flip a coin to choose the left–right order of children of v;
        Inorder-Traversal(left child of v, σ, a, b);
        append the height of v to a;
        Inorder-Traversal(right child of v, σ, a, b);
    else
        append the label of v to σ;
        append the height of v to b;
    end
end
Function Reconstruct-Tree(linked list of nodes s of length m) is
    let i be the index of the highest internal node in s;
    if i = 2 then
        set the left child of s_i to s_1;
    else
        set the left child of s_i to Reconstruct-Tree(s_{1,...,i-1});
        if the height of s equals the height of its left child then reject the proposal;
    end
    if i = m − 1 then
        set the right child of s_i to s_m;
    else
        set the right child of s_i to Reconstruct-Tree(s_{i+1,...,m});
        if the height of s equals the height of its left child then reject the proposal;
    end
    return s_i;
end
let σ, a, a*, b and s be empty linked lists;
Inorder-Traversal(the root node of T, σ, a, b);
/* j points to the true bifurcation MRCA height to change and is also the length of X_L */
let j be a random integer for which a_j ≠ b_j and a_j ≠ b_{j+1};
let X_l be the sampled alleles which map to X_L = σ_{1,...,j};
let X_r be the sampled alleles which map to X_R = σ_{j+1,...,n};
let upperBound be ∞ or, if it is available, O;
for x ∈ g ∈ G do
    let x_l be the sampled alleles descended from the left child branch of x;
    let x_r be the sampled alleles descended from the right child branch of x;
    if (x_r ∩ X_r ≠ ∅ and x_l ∩ X_l ≠ ∅) or (x_r ∩ X_l ≠ ∅ and x_l ∩ X_r ≠ ∅) then
        set upperBound to the height of x if this lowers upperBound
    end
end
let lowerBound be the larger of a_j or a_{j+1};
foreach e ∈ a do append e to a*;
let h ~ uniform(a_j − δ, a_j + δ);
reflect h against lowerBound and upperBound until lowerBound < h < upperBound;
change a*_j to equal h;
for i = 1 to n do
    append a new leaf node with the label σ_i and height b_i to s;
    if i < n then
        append a new internal node of height a*_i to s;
    end
end
initialize and return T* using Reconstruct-Tree(s) as the root;
```

#### Correctness of tree topologies and node heights

There are three relevant sets of operators to check: BEAST 2 operators, StarBEAST2 operators including `TREE SLIDE`, and Sampled Ancestor operators. The FBD-MSC implementation in StarBEAST2, employed in our empirical analyses, uses operators from all three sets. One way to verify these operators is to look at manageable, small-size data sets for which we can compute (true) phylogenetic expectations, for both topology and divergence times. Such an approach was used to validate the implementation of FBD-concatenation in BEAST 2 [2], which we follow here as well.

We replicated the model parameters used for the original validation of operators for FBD-concatenation in BEAST 2 [2]. These are:

- Birth-rate, $\lambda = 2.0$;
- Death-rate, $d = 1.0$;
- Sampling-rate, $\psi = 0.5$;
- Removal probability, $r = 0.9$;
- Origin time, $\mathcal{O} \sim \mathrm{Unif}(0, 1000)$.

The number of species is fixed at three, and the sampling times of those species are fixed at $y_1 = 2.0$, $y_2 = 1.0$, and $y_3 = 0.0$. A maximum of three parameters are sampled (the origin time $\mathcal{O}$ and two bifurcation times). If the species serially sampled at $y_1$ or $y_2$ are sampled ancestors, then there is either none or one bifurcation time to estimate. There are eight possible topologies given one extant and two fossil species, and we used previously calculated frequencies [2] for their true probabilities absent any data (Table S1).

| Topology | Newick string | Probability (%) |
|---|---|---|
| T1 | ((3,2),1) | 77.8327 |
| T2 | ((3,2)1) | 7.8642 |
| T3 | ((3)2,1) | 3.8657 |
| T4 | (3,(2,1)) | 4.3189 |
| T5 | ((3,1),2) | 4.3189 |
| T6 | (((3)2)1) | 0.4135 |
| T7 | (3,(2)1) | 0.6930 |
| T8 | ((3)1,2) | 0.6930 |

Table S1: Possible three-taxon topologies and corresponding probabilities for the FBD model and parameter values we specified. ((3,2)1), for example, indicates "1" is a sampled ancestor of (3,2); (((3)2)1) indicates "1" and "2" are sampled ancestors of "3", with "2" being younger.

We note that the coalescent probability densities are conditioned on the species tree (the FBD-MSC model is hierarchical) and therefore, absent any data, the addition of one or more gene trees to the model should not change the distribution of species tree topologies or times.

Keeping the parameters from the model described above constant, we set up six different BEAST 2 analyses, each with a different combination of operators. The logic here is to use operators in isolation in order to detect issues that might be present in a few, but not all of them:

- "SA": All the Sampled Ancestor operators (see "Previously developed MCMC operators") applied to a species tree. No gene trees were part of the model and StarBEAST2 was not activated.
- "UpDown": The same as "SA", but with the `UpDown` BEAST 2 operator enabled for the species tree.
- "MSC": StarBEAST2 with one gene tree, the same operators as "UpDown" for the species tree, and the BEAST 2 operators (again see "Previously developed MCMC operators") listed above for the gene tree. No StarBEAST2 operators were enabled.
- "Coordinated": The same as "MSC" but `CoordinatedExchange` and `CoordinatedUniform` were enabled.
- "NodeReheight2": The same as "MSC" but the `NodeReheight2` operator was enabled.
- "Full": The same as "MSC" but `CoordinatedExchange`, `CoordinatedUniform` and `NodeReheight2` were all enabled.

For each configuration, we ran 100 replicate MCMC chains with different random seeds. Each chain was run for 2.5 million states with a burn-in of 0.5 million states, sampling once every 100 states. Therefore 20,000 post-burn-in posterior samples were collected per replicate chain.

##### Tree topology correctness

Given $n$ independent trials with probability of success $p$, the number of "successes" should follow the binomial distribution parameterized by $n$ and $p$. Therefore for each topology, the range defined by the 2.5-th and 97.5-th percentile of a binomial distribution with the parameter values $n =$ effective sample size (ESS), and $p =$ the true probability of a topology, after normalization by the effective sample size, should contain the estimated probability of that topology 95% of the time.

For each MCMC chain we calculated the estimated probability of each topology as the number of observations of that topology divided by the total number of samples. We also calculated the autocorrelation time for each topology in each chain by coding samples containing that topology as 1 and all other samples as 0, then using Sokal's adaptive truncated periodogram estimator [8] as implemented in the `integrated_autocorrelation_time` function from `pymcmcstat`. The ESS for each topology in a chain is simply the raw number of samples divided by the estimated autocorrelation time.

For each topology in each configuration, we counted the number of replicate chains for which the estimated probability of that topology was within the 95% interval.

##### Tree node height correctness

Fundamentally, MCMC is an algorithm for performing difficult high dimensional integrations, e.g., calculating the marginal probability distribution of the height of a node in a phylogenetic tree. For the species tree topologies in this analysis we are estimating a maximum of three real parameters, so it is not necessary to use MCMC; we can use an alternative implementation of the FBD to calculate the marginal probability distributions of those real parameters. If both our MCMC and non-MCMC implementations are correct, the estimated distributions should be identical (up to some small error term).

We re-implemented the FBD model for the T1 species tree topology – ((3,2),1) – given our model. This implementation was written in Python 3 using the `SciPy` library, which in turn calls the `QUADPACK` library to integrate over any interval using adaptive or non-adaptive quadrature.

For each real parameter (the origin time $\mathcal{O}$, the root node height $x_1$, and the non-root internal node height $x_2$), separately for each replicate MCMC chain, we calculated probability masses conditioned on the T1 species tree topology. Masses were calculated for 100 equal width intervals (0.1 time units each) between 0 and 10. For each interval we recalculated the same probability masses using our Python implementation.

#### Validating the full model

We simulated parameter values (Fig. S4, Table S2) from their (hyper-)priors, and then simulated discrete morphological traits and molecular alignments conditioning on those values. Sampling distributions are listed in table S3. Species trees were sampled from the FBD model, and given those species trees, gene trees were sampled from the MSC model.

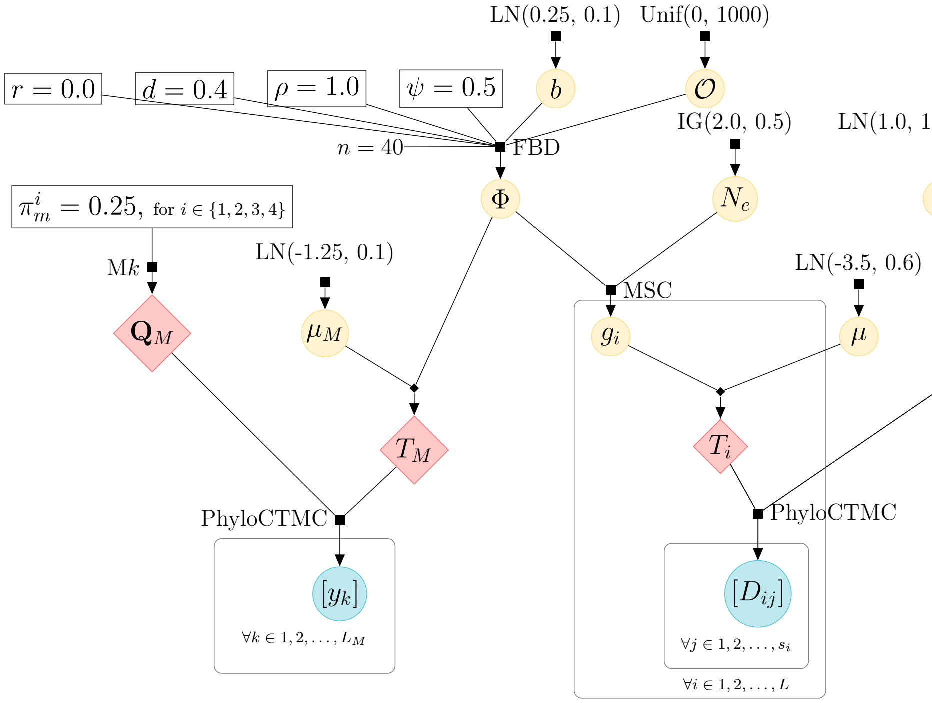


Figure S4: Full probabilistic graphical model used in the well-calibrated validation. Filled black squares and diamonds represent sampling distributions and deterministic functions, respectively. Yellow and blue filled circles denote random variables and data, respectively. Symbols enclosed in unfilled rectangles correspond to constants. In our model, $L = 4$, $s_i = 500$, $L_M = 100$, and species trees have $n$ extant species. $\mathbf{Q}$ and $\mathbf{Q}_M$ are the instantaneous rate matrices determined by the molecular and morphological substitution models given their parameters. $T_i$ and $T_M$ are the i-th molecular tree (out of 4) and the morphological tree respectively, both in units of expected substitutions per character. Remaining variable symbols and sampling distributions are described in tables S3 and S2.

We constrained our species tree simulations to species trees with $n = 40$ extant species (the number of extinct and sampled ancestor species were still variable; Fig. S9). Here, simulations are done backwards in time, and stop when the origin is reached (there is no explicit prior on the origin time, $\mathcal{O}$, i.e., the total tree height). Species trees much smaller or much larger than this can be either uninteresting problems (in the case of very small trees, certain parameters are unidentifiable), too computationally expensive (i.e., if trees are too large, MCMC chains do not converge in a reasonable amount of time), or unrealistic (very few clades have large data sets of real genomic, fossil and morphological data). Conditioning on 40 extant species is thus advantageous in that it allows us to keep trees at both realistic and useful sizes for model validation. A drawback, however, is that this conditioning creates must create a a mismatch between the FBD

Table S2: Summary of sampling (prior) distributions used in well-calibrated validation study. Distribution descriptions can be found in Table S3.

| Parameter | Description | Sampling distribution |
|---|---|---|
| | **Fossilized birth-death (FBD) parameters** | |
| $b$ | Birth rate | LN(0.25, 0.1) |
| $d$ | Death rate | 0.4 (fixed) |
| $\psi$ | Sampling rate | 0.5 (fixed) |
| $\rho$ | Extant sampling proportion | 1.0 (fixed) |
| $r$ | Removal probability | 0.0 (fixed) |
| | **Multispecies coalescent (MSC) parameters** | |
| $\Phi$ | (see below) | |
| $N_e$ | Effective population sizes once scaled by generation time | IG(2.0, 0.5) |
| | **Phylogenetic trees** | |
| $\Phi$ | Species tree | FBD$(b, d, \psi, \rho, r)$ |
| $g_i$ | i-th gene tree | MSC$(\Phi, N_e)$ |
| | **Molecular substitution model (HKY) parameters** | |
| $\kappa$ | Transition:transversion ratio | LN(1.0, 1.25) |
| $\pi$ | Nucleotide equilibrium frequencies | Dir(1,1,1,1) |
| | **Morphological substitution model (Mk) parameters** | |
| $\pi_m$ | State equilibrium frequencies | $\pi_m = \{0.25, 0.25, 0.25, 0.25\}$ (fixed) |
| | **Clock model parameters** | |
| $\mu$ | Global molecular rate | LN(-3.5, 0.6) |
| $\mu_M$ | Global morphological rate | LN(-1.25, 0.1) |

Table S3: Sampling distributions we used in our empirical and validation analyses.

| Sampling distribution | Description |
|---|---|
| LN($\mu$, $\sigma$) | Log-normal distribution with log-space mean $\mu$ and log-space standard deviation $\sigma$ |
| Dir($\cdot$) | Dirichlet distribution |
| U($a, b$) | Uniform distribution for interval $[a, b]$ |
| Exp($\lambda$) | Exponential distribution with rate $\lambda$ |
| Beta($\alpha$, $\beta$) | Beta distribution with shape parameters $\alpha$ and $\beta$ |
| IG($\alpha$, $\mu$) | Inverse gamma distribution with shape parameter $\alpha$ and mean $\mu$ |

model used in simulation (here) and the one used in inference in the main manuscript, which does not condition on the number of extant species. Given our validation results, this model misspecification does not seem serious – but if anything, it makes our validation results conservative.

Alternatively, species trees can be simulated forward in time by assigning a prior on origin times, and stopping simulations at those times. While in principle this strategy does not cause model misspecification, species trees will then exhibit a wide range of tip counts, going from anywhere between one species to several hundred. This is a known problem for the validation of birth-death models [2], one solution for which is to then filter out trees that have too few or too many species for the reasons mentioned above. Excluding simulated trees, however, in turn introduces a model misspecification as well: trees that are thrown away are effectively being set prior probabilities of zero, but during inference those probabilities will be non-zero. We chose to not filter trees out, and instead conditioned our simulations on 40 extant species as described above.

We ran a total of 100 MCMC chains (one per simulated data set) for 50 million steps each, discarding the first 10% as burn-in. All but three converged (i.e., all but three had a prior, posterior, or likelihood with an effective sample size greater than 100); those three were excluded from further analysis.

Table S4: Software packages employed in well-calibrated validation analysis.

| Package | Type | Version | Use | Reference |
|---|---|---|---|---|
| `FossilSim` | R package | 2.1.1 | FBD tree simulation | [9] |
| `phangorn` | R package | 2.5.5 | Discrete trait simulation | [10] |
| `phytools` | R package | 0.7.47 | Phylogenetic tree parsing and visualisation | [11] |
| `Unreleased BEAST 2 package` | Java code | N/A | Gene tree simulation under the MSC and molecular alignment simulation | N/A |

#### Model and priors for the Caninae analyses

The FBD-MSC model used to infer a posterior distribution of Caninae species trees $\Phi$ and other parameters is presented as a graphical model in Fig. S5. The detailed justifications for the prior distributions we chose can be found in the text. Distributions are specified as functions (Table S3).

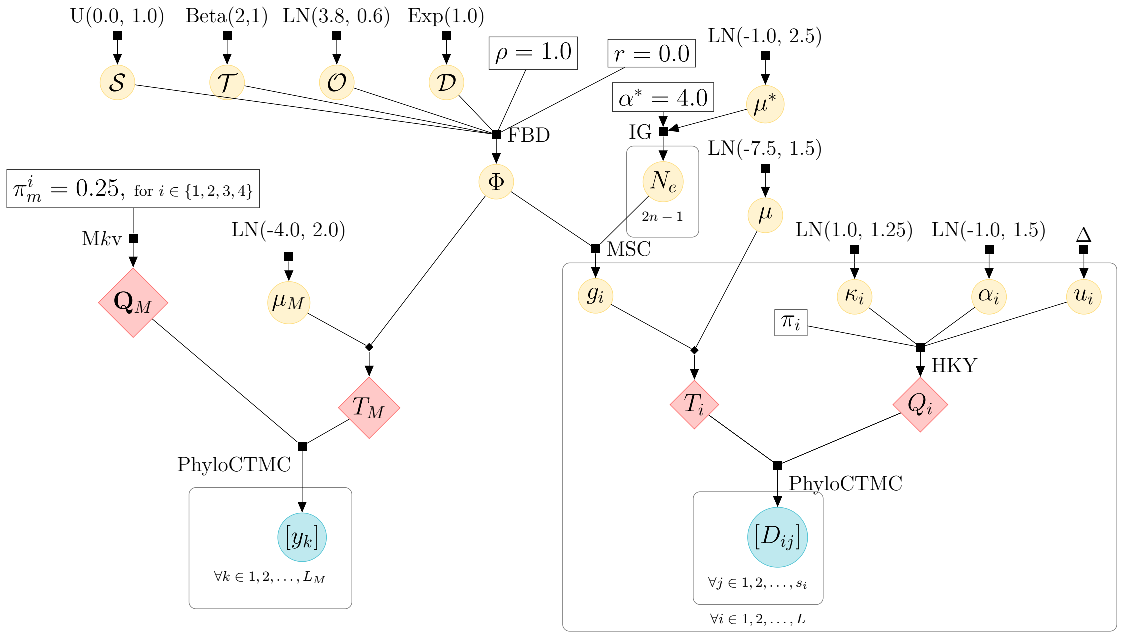


Figure S5: Full probabilistic graphical model used in the FBD-MSC analysis of Caninae. Filled black squares and diamonds represent sampling distributions and deterministic functions, respectively. Yellow and blue filled circles denote random variables and data, respectively. Symbols enclosed in unfilled squares correspond to constants. We use $\alpha^*$ $\mu^*$ for the shape and mean of the inverse gamma prior on $N_e$, to distinguish those parameters from the rate variation and clock rate parameters $\alpha$ and $\mu$. We use $\Delta$ to represent the constrained uniform prior on relative substitution rates. In our model, $L$, $s_i$, $L_M$, $n$, and $\pi_i$ come directly from the data set. $\mathbf{Q}$ and $\mathbf{Q}_M$ are the instantaneous rate matrices determined by the molecular and morphological substitution models given their parameters. $T_i$ and $T_M$ are the i-th molecular tree and the morphological tree respectively, both in units of expected substitutions per site. Other symbols are explained in the main text.

###### Species tree parameters and priors

For the Caninae analysis we used the alternative FBD parameterization of net diversification $\mathcal{D} = b - d$, turnover $\mathcal{R} = d/b$ and fossil sampling proportion $\psi/\psi+d$ [12], where $b$, $d$ and $\psi$ refer to the birth, death and sampling rates respectively.

**Origin,** $\mathcal{O} \sim$ LN$(3.8, 0.6)$. We chose this prior based on the oldest carnivoran fossils (not in our data sets), which appear in the Paleocene, around 56–66 mya [13, 14]. Our prior has an 50% equal-tailed interval (ETI) spanning 30 to 67 million years ago (mya), a wide range of origin values with non-negligible prior probabilities. Note that the oldest fossil in our data set is approximately 30 million years old.

**Net diversification,** $\mathcal{D} \sim$ Exp$(1)$. We reasoned that branches in a tree of mammalian species should last a period of time on the order of millions of years, and a net diversification rate of 1.0 is expected to lead to one bifurcation every million years. Because the duration of a species can vary considerably, however, we chose an exponential prior. Our prior assigns higher densities to diversification values closer to zero (implying longer intervals between speciation events), but still has a long tail beyond the mean to allow for shorter internal branches (as one might observe in more radiation-like evolutionary scenarios). The 95%-ETI spans 0.025 to 3.68, which allows for waiting times between speciation events of roughly 0.3 to 40 million years.

**Turnover,** $\mathcal{T} \sim$ Beta$(2, 1)$. Extinction must be common among Canidae since the two other subfamilies are entirely extinct, as well as for other vertebrates given their extensive fossil record. We thus chose a prior for the turnover fraction that is monotonically increasing from 0.0 to 1.0, as values close to 0.0 would also imply very low death rates incompatible with the ubiquitous level of extinction seen in the real world.

**Fossil sampling proportion,** $\mathcal{S} \sim$ U$(0, 1)$. When this value is greater than half, sampling is more frequent than extinction. When it is less than half, it is less frequent than extinction. Given that this parameter has natural bounds of 0.0 and 1.0, and because we do not have enough prior knowledge about what its value should be, we employed a uniform prior.

**Extant sampling proportion,** $\rho = 1$ . This value implies complete sampling of extant species. While the taxonomy of Caninae is fluid, the study we adapted the trait matrix from attempted to represent the extant interspecies diversity of the subfamily by including all unproblematic extant species and appropriate representatives of the dog/grey wolf/coyote species complex [15]. The only exception is the recent splitting of *Canis aureus* and *C. anthus* [16]. We therefore believe complete sampling of extant species to be a reasonable approximation.

**Removal probability,** $r = 0$ . Positive values imply that sampling a tip can lead to the extinction of that lineage. Since this connection does not exist for fossils, we fix removal at zero.

###### Substitution model parameters and priors

All substitution model parameters are estimated individually for every $i$-th locus.

**Transition:transversion ratio,** $\kappa_i \sim$ LN$(1, 1.25)$. Previous studies of this ratio for mammalian nuclear genes, a category most of the genes in our data set belong to, found the average rate ratio $\kappa$ is around 3 and no genes with rate ratios below 1 or above 6 [17, 18]. We kept the default BEAST prior on $\kappa_i$ because its 50% credible interval spans approximately 1 to 6.

**Gamma-rate variation shape,** $\alpha_i \sim$ LN$(-1, 1.5)$. This parameter is generally estimated to be less than 1 [19, 20]. We chose this prior on the gamma-rate variation shape $\alpha_i$ because it has a 50% ETI spanning 0.13 to 1, encompassing the values observed on those studies, but a much wider 95% ETI spanning 0.02 to 7. This leaves room for values substantially higher than 1 in case some loci had relatively little rate variation.

**Nucleotide equilibrium frequencies,** $\pi_i$ fixed at (observed) empirical frequencies. Given a multiple sequence alignment of sufficient size, the observed (empirical) base frequencies should be a good estimate of $\pi_i$.

**State equilibrium frequencies,** $\pi_m^i$ fixed at equal frequencies. Given the limited information content for each trait we used equal frequencies.

**Relative substitution rates,** $u$ constrained to a weighted (by the number of sites in each locus) average of 1. This is achieved by exclusively using the "Delta exchange" operator to change this parameter, and helpfully causes the global clock rate to exactly equal expected substitutions per site per time unit. The prior is uniform but not improper thanks to the constraint, and may be formalized as $\begin{cases} 1/Z & \text{if} \sum_i u_i s_i = \sum_i s_i \\ 0 & \text{if} \sum_i u_i s_i \neq \sum_i s_i \end{cases}$ where $Z$ is the normalizing constant for this uniform distribution.

**Clock model parameters and priors**

**Global (absolute) molecular clock rate,** $\mu \sim$ LN$(-7.5, 1.5)$. Mammalian nuclear coding sequence clock rates are on the order of $1 \times 10^{-3}$ substitutions per site per year [21]. We therefore chose a log-normal prior with a similar mean of about $1.7 \times 10^{-3}$. The 95% ETI of this distribution spans $3 \times 10^{-5}$ to $1 \times 10^{-2}$, making it a good fit for our expectations that the clock rate will be within an order of magnitude of $1 \times 10^{-3}$.

**Global morphological clock rate,** $\mu_M \sim$ LN$(-4, 2)$. Traditionally, taxonomists have identified and coded characters that are both variable and synapomorphic (defining a monophyletic group). If we consider a tree with $n = 100$ extant taxa and such types of characters, then there will be $n - 1$ internal branches that could undergo a character-state transition. A rate that induces transitions on all branches is unlikely, as that would mean each species would have its own state and the character would be useless for recovering the tree (and thus not scored in the first place) – i.e., the morphological alignment would be saturated. If each internal branch is on the order of 1 million years, then we would expect rates to be much lower than 1 substitution per character per million years; more specifically if characters generally consist of synapomorphies, then the median rate should revolve around $\frac{1}{(n-1)}$. We note, however, that because the morphological rate under the Mkv model takes into account invariable characters (i.e., ascertainment bias is controlled for), our prior on rate values should be skewed toward values below the median. The prior we chose observes these expectations, having a median of 0.01, and a 95%-ETI interval spanning $4 \times 10^{-4}$ and 0.9.

**Demographic parameters and priors**

**Population sizes,** $N_e \sim \text{IG}(4, \textit{estimated})$. An inverse gamma prior on population sizes is used because it can be analytically integrated over, improving computational performance. Prior to StarBEAST2, *BEAST was widely used for a decade and implemented a fixed, hard-coded gamma prior on population sizes with a shape of 2. We chose 4 for the value of $\alpha$, because its coefficient of variation is $\frac{\sqrt{2}}{2}$ is identical to the coefficient of variation of the tried and tested *BEAST demographic model. We chose to estimate the mean of this distribution as the population scale (see below).

**Population scale** $\sim$ LN(-1, 2.5). This is the hyperprior for the mean of the population sizes prior. StarBEAST2 population sizes are in the units $N_e g$ where $N_e$ is the population size of a Wright-Fisher population and $g$ is the generation time in millions of years. In order to choose an adequate prior for this parameter, note that nucleotide diversity is an estimator of $\theta = 4N\mu$, where $\mu$ is substitutions per site per generation. $\theta$ in vertebrates can range from between $1 \times 10^{-4}$ and $1.5 \times 10^{-2}$ [22]. Assuming $\mu = 1 \times 10^{-3}$ substitutions per site per million years, that converts to population sizes in $N_e g$ from as low as 0.025 to as high as 3.75. Accordingly, we used a prior with a 95% ETI encompassing an order of magnitude beyond those numbers in both directions.

#### Rate and divergence time estimates depend not only on the MSC, but also on the dating method

In this section, we address the valid question of whether tip dating plays any role in correcting the estimation of molecular rates and divergence times, or whether better estimates can be obtained using the MSC model alone. If the MSC is the only model component that matters for the purpose outlined above, then utilizing another dating method should yield similar results to those from the FBD-MSC model. Otherwise, one must conclude that tip dating (via the FBD) must matter, at least in some situations (such as the case of Caninae evolution).

In order to answer this question, we carried out a node dating analysis of the same molecular and morphological data, under a model referred to as the birth-death-MSC (BD-MSC). As mentioned in the main text and above, because node dating is ad hoc in nature, our model configuration is just one of an infinite number of possible node-dating models.

Here, one could rightfully object that whatever results we report next are contingent upon the chosen calibration scheme. Our goal, however, is not to demonstrate that the FBD-MSC will **always** lead to different results than node dating, but instead that it **can**. This conclusion should in fact *follow trivially* from the understanding that there is as myriad of ways one could do node dating, i.e., that tip dating must occasionally lead to different results when compared to other tree priors in conjunction with the MSC. We nonetheless pick one possible calibration scheme to test the hypothesis that the MSC does not always do the “heavy-lifiting” by itself.

Finally, we note that we do not advocate for node dating – there are multiple critical reasons why it should be avoided, as mentioned in the main text and covered in detail in [23–25] – and instead recommend tip-dating (e.g., using the FBD) whenever possible.

##### BD-MSC configuration

Under the BD-MSC model, we included only extant species and the recently extinct *Dusicyon australis* as a calibration specimen. We calibrated the split of the North American and old world (including *Canis*, *Lycaon*, *Cuon* and *Lupulella*) and South American (including *Lycalopex*, *Dusicyon*, *Cerdocyon*, *Atelocynus*, *Chrysocyon* and *Speothos*) radiations within Canini, reusing a previously developed calibration for the age of the North American radiation [26]. This calibration is an exponential distribution with a mean of $1.\overline{6}$ million years, offset by 5 million years. Unlike for FBD models, there is no removal probability or fossil sampling proportion, and we conditioned the species tree on the root instead of the origin. For all other parameters of the model, including the of net diversification, turnover and extant sampling proportions of the birth-death model, we used the same values or prior distributions as for FBD-MSC.

The BD-MSC analysis consisted of four independent $2^{26}$-generation (roughly 67 million) MCMC chains. Again, chains were joined after discarding the first 64 samples (roughly 1.6%) from each chain as burn-in based on the manual inspection of MCMC traces in Tracer 1.7 [27].

##### Computational performance

The lowest effective sample sizes was observed for the gamma distribution shape. Performance under the BD-MSC model is summarized in Table S5.

Table S5: Computational performance of Caninae analyses for BD-MSC.

| Model | Combined CPU hours | Min. ESS | ESS per hour | Parameter with min. ESS |
|---|---|---|---|---|
| BD-MSC | 16 | 488 | 30.5 | *PNOC* gamma shape |

##### BD-MSC results

Posterior estimates under the BD-MSC model are summarized in Table S6. Node dating under the BD-MSC model yielded a mean molecular rate posterior estimate of $7.3 \times 10^{-4}$, in between those from the two tip-dating models, with a more spread out posterior distribution (Fig. S6a).

We show for the first time that leveraging serially sampled data in conjunction with the MSC (i.e., the FBD-MSC model) can significantly improve rate estimates and better inform node age inference relative to node dating. This is evidenced by node age estimates under the BD-MSC model being more uncertain, as well as larger, undercorrected molecular rate mean posteriors when compared to those obtained when using the FBD-MSC.

Table S6: BD-MSC parameter estimates. All values are posterior mean estimates followed in brackets by 95% highest posterior densities. $^*$per million years. $^{**}$the mean of the inverse gamma distribution fit to per-branch $N_e g$ values, which are effective population sizes $N_e$ scaled by generation time in millions of years $g$.

| Parameter | BD-MSC |
|---|---|
| Molecular clock rate ($\times 10^{-3}$) | 0.73 (0.48–0.93) |
| Morphological clock rate ($\times 10^{-2}$) | 3.95 (2.62–5.17) |
| Mean population size$^{**}$ | 1.34 (0.76–2.00) |
| Diversification rate ($\lambda - \mu$) | 0.25 (0.10–0.40) |
| Turnover ($\mu \div \lambda$) | 0.35 (0.04–0.69) |
| Sampling proportion ($\psi \div (\psi + \mu)$) | NA |

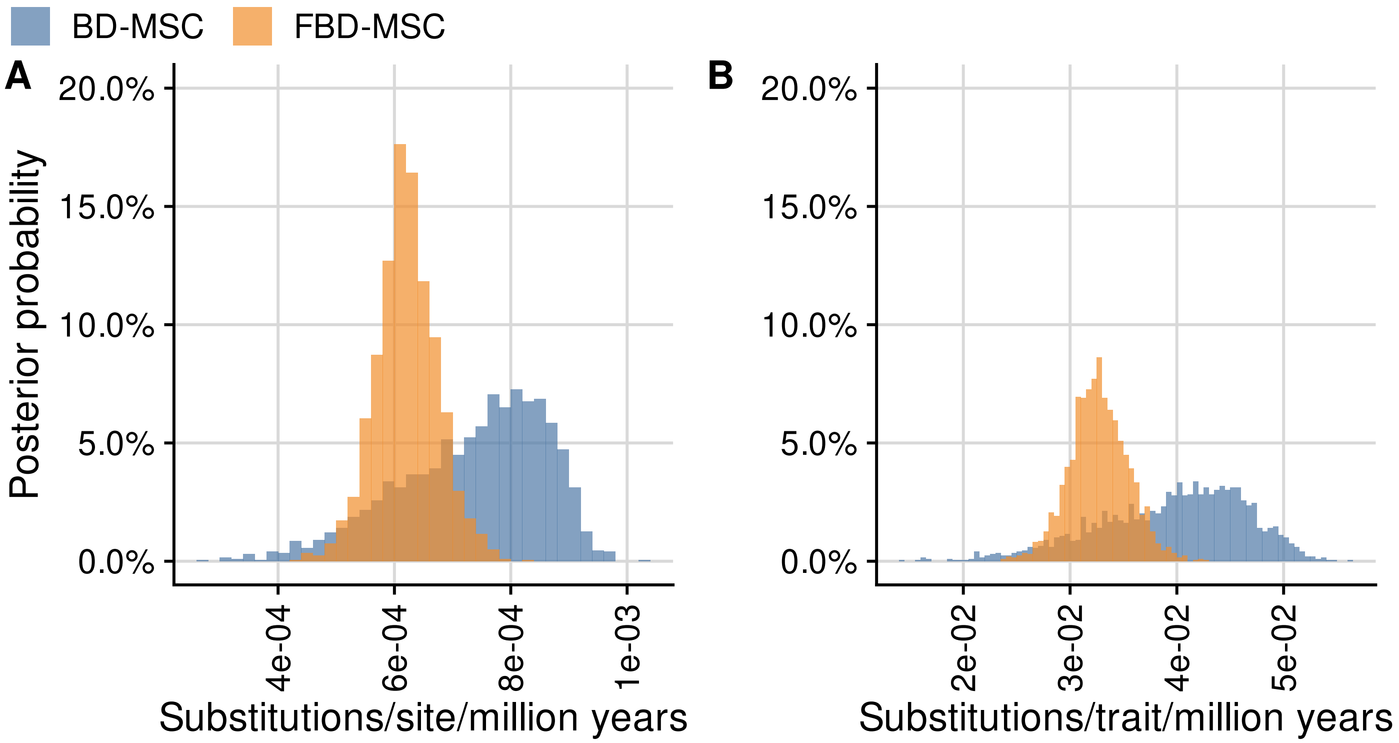


Figure S6: Posterior distribution of clock rates using MSC models. Posterior probabilities of molecular clock rates (A) and morphological clock rates (B) were calculated using bin widths of $2 \times 10^{-5}$ and $5 \times 10^{-4}$ respectively.

Morphological clock rate estimates under the BD-MSC model were also uncertain, to an even greater extent – perhaps unsurprisingly given that these estimates came from a smaller number of data points (we only use living species) – as indicated by a much wider posterior distribution centered at larger morphological clock rate values (Fig. S6b).

Finally, the BD-MSC model produced node log-age estimates in agreement with FBD-MSC, albeit more uncertain, as reflected by wider 95% HPDs (Fig. S7 and Fig. S8).

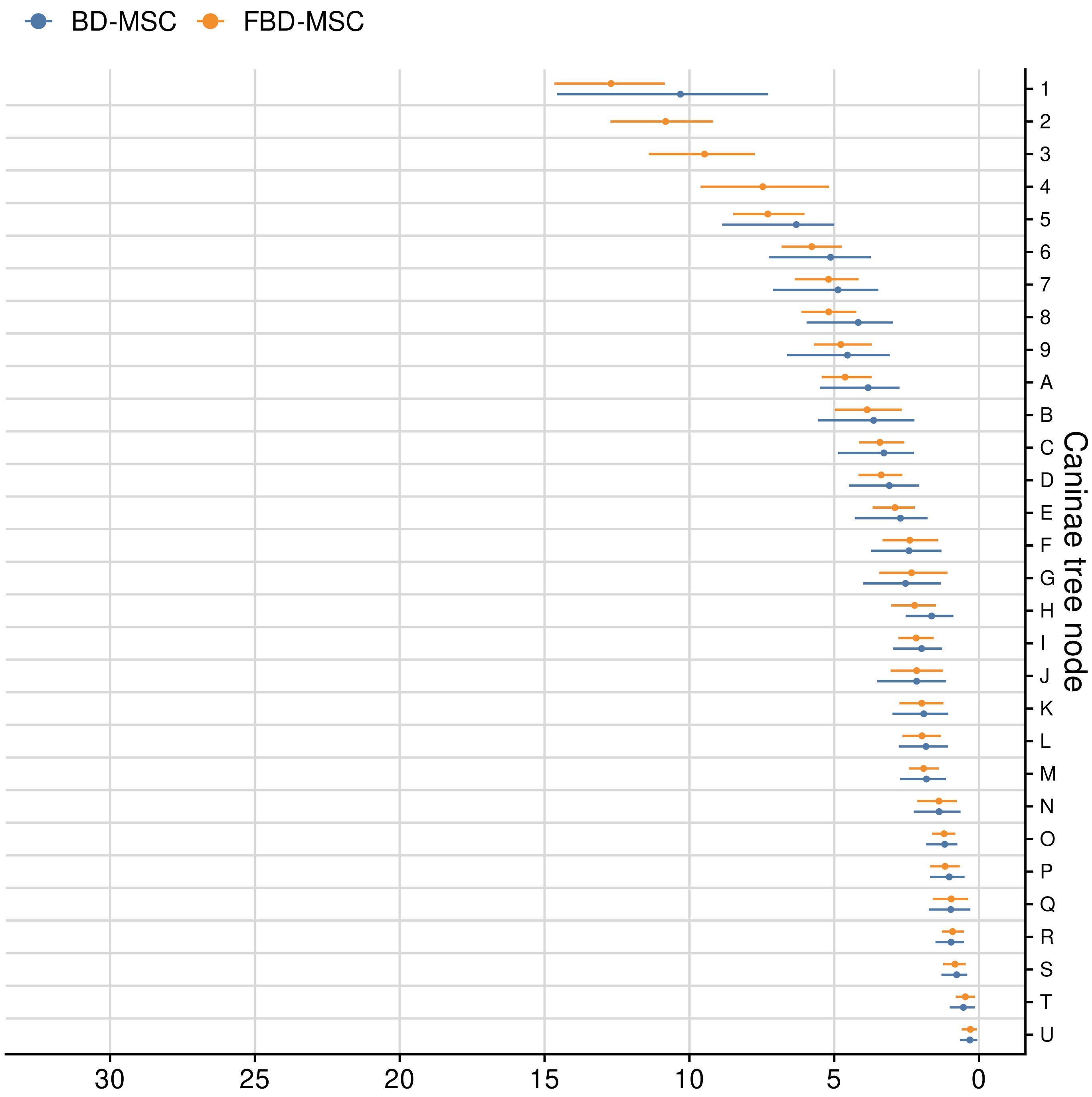


Figure S7: Tempo of Caninae evolution. Crown ages estimated by fossilized birth-death with multispecies coalescent (FBD-MSC) and with a birth-death and multispecies coalescent model (BD-MSC) (a), compared with lineages-through-time (LTT) curves including extinct lineages (b). Posterior mean internal node ages (solid circles) with 95% highest posterior density (HPD) intervals are estimated from samples where that clade is present after pruning all morphology-only taxa. 95% highest posterior density (HPD) intervals calculated for each step are shown as ribbons.

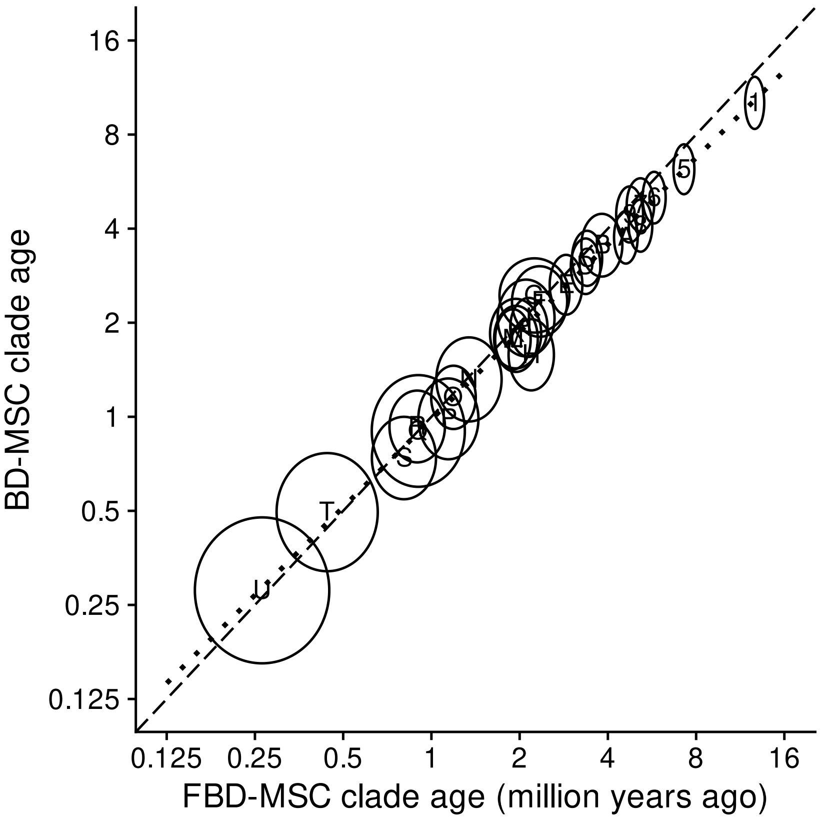


Figure S8: Correlation between log-node heights from the posterior distributions of species trees pruned of morphology-only taxa. Internal nodes from the pruned FBD-MSC MCC tree are drawn as ellipses centered on the mean estimate of the log-node height for both methods. The width and height of each ellipse corresponds to the standard deviation of the log-node heights for FBD-MSC and BD-MSC respectively. The dashed black line shows the 1:1 line along which estimates are equal, and the solid blue line is the quadratic line of best fit.

### Additional figures and tables

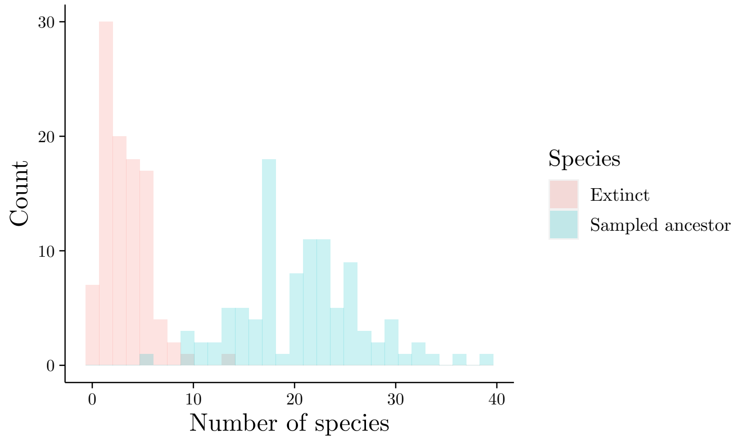


Figure S9: Distribution of species counts in simulated species trees during well-calibration study. Extinct species correspond to fossil tips.

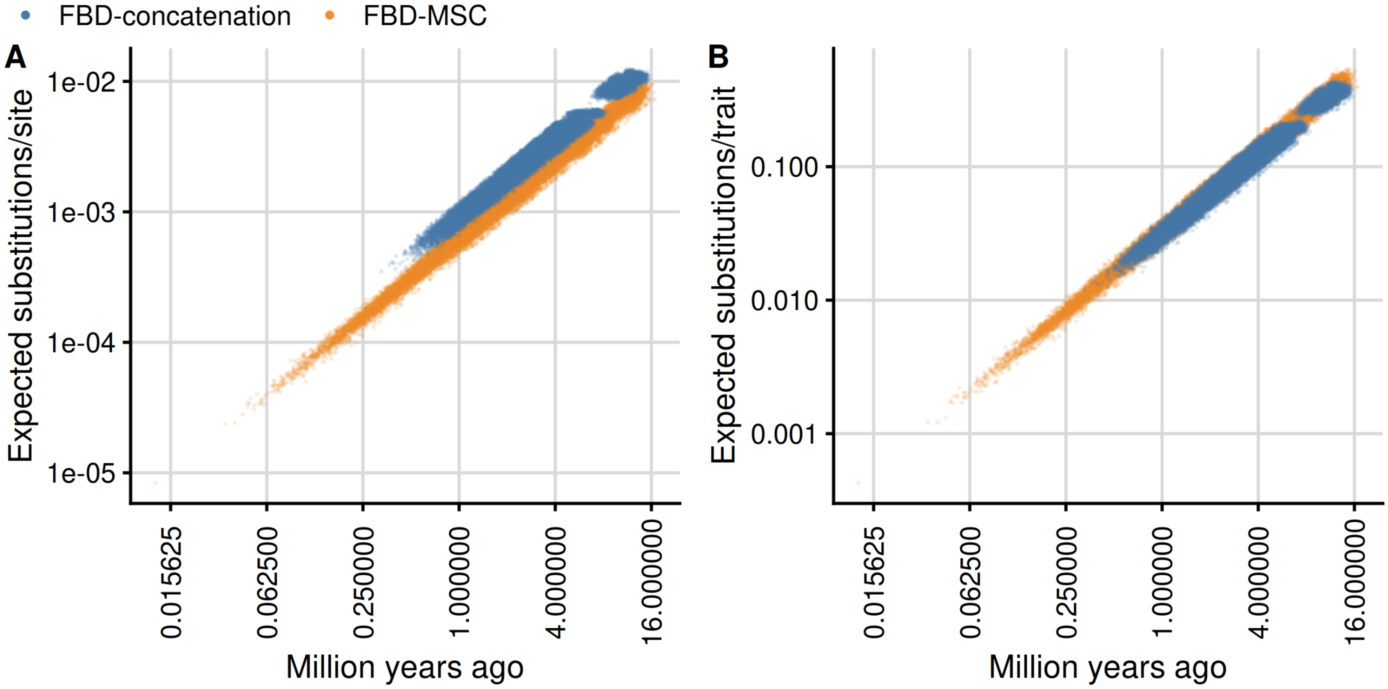


Figure S10: The age of every node in every posterior sample in expected substitutions/site is a function of the age in Ma and the molecular clock rate for that sample (C), and its age in expected substitutions/trait is a function of its age and the morphological clock rate for that sample (D).

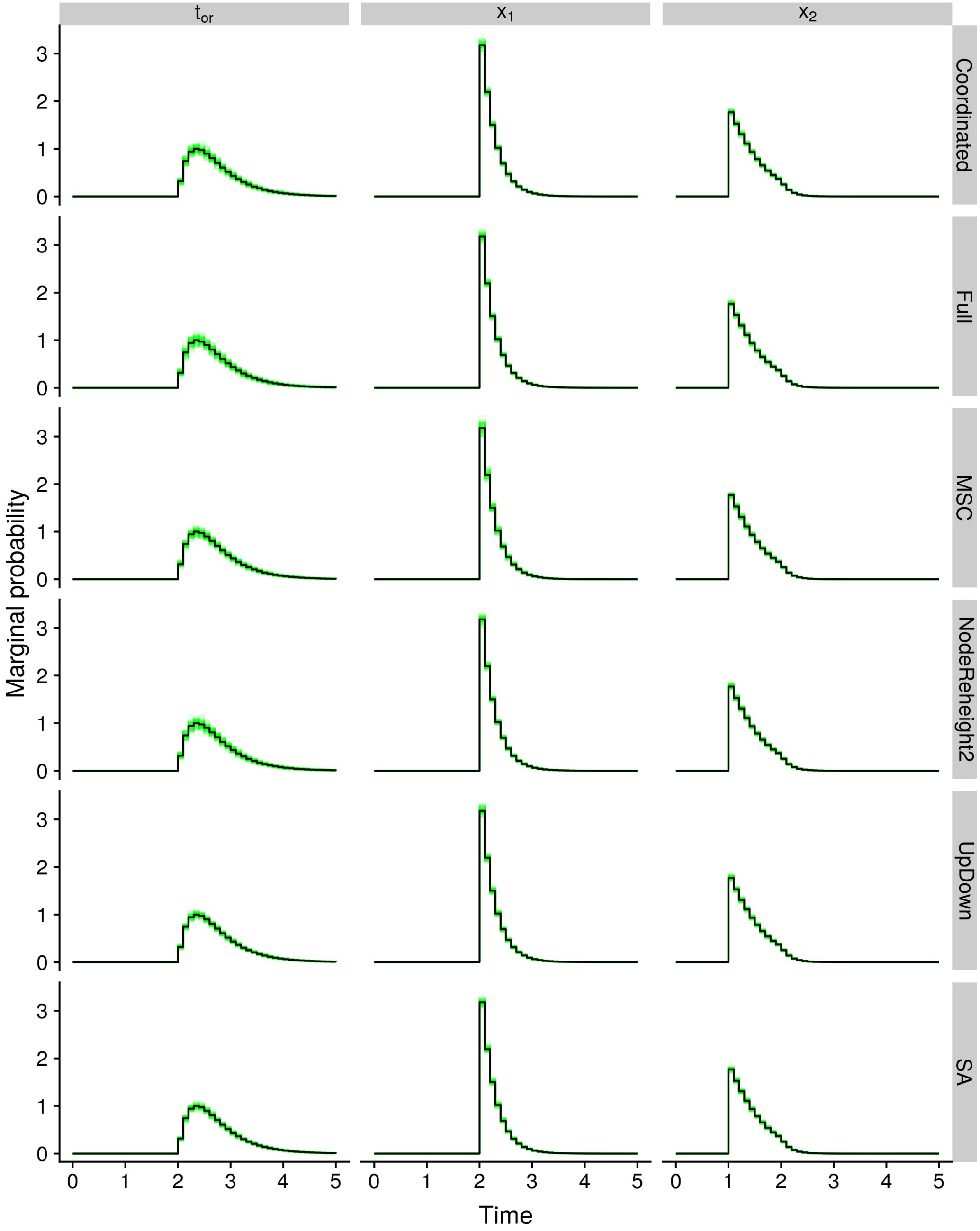


Figure S11: Marginal prior distributions of the origin ($t_{or}$) and speciation ($x_1$, $x_2$) times. Distributions were computed using quadrature (black histograms), or inferred using StarBEAST2 MCMC with different sets of operators enabled (green histograms). Plots were truncated at time = 5 since marginal probabilities are so low from that point.

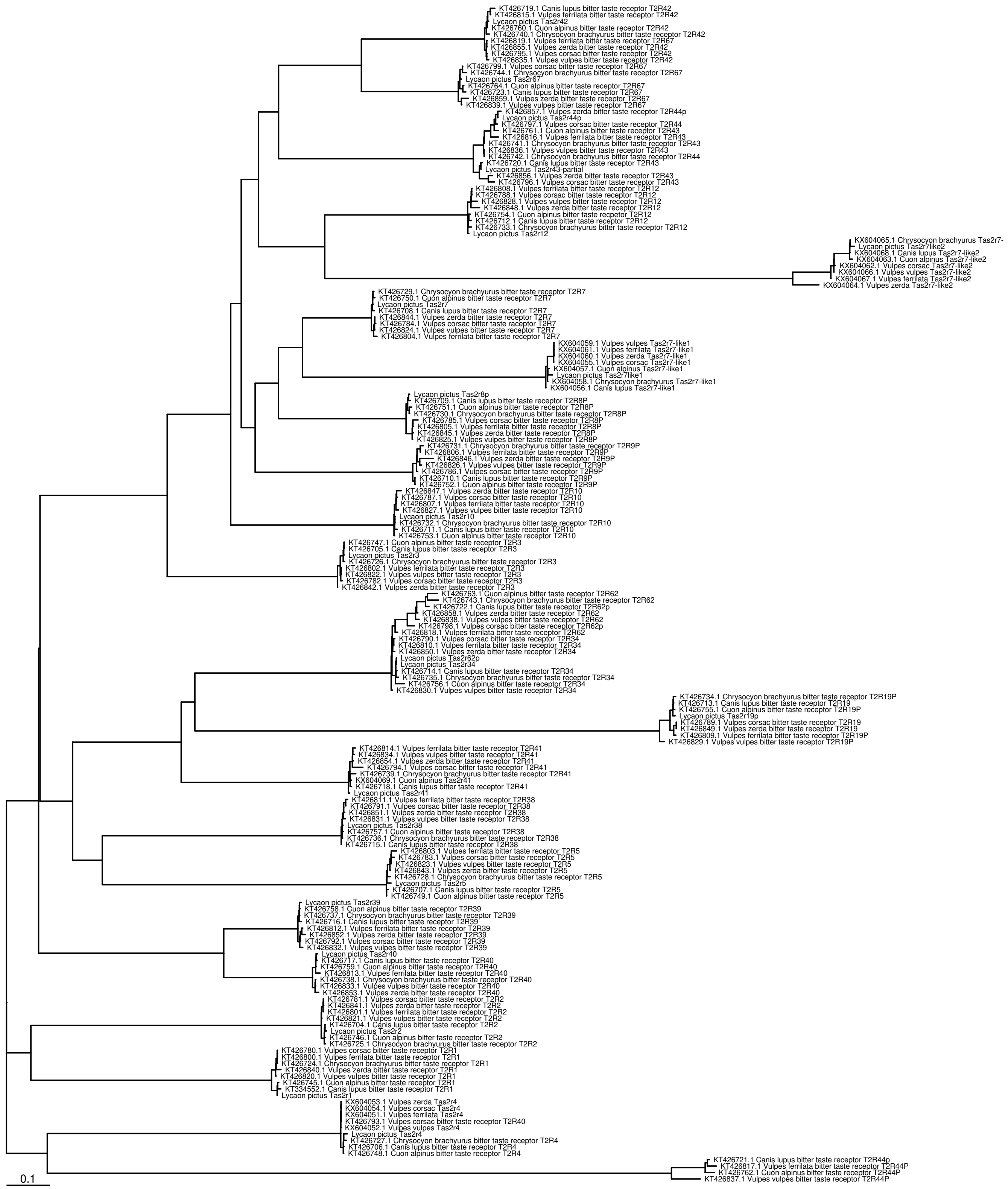


Figure S12: Tree of taste 2 receptor (*Tas2r*) genes. Labels for non-*Lycaon* sequences, which are not available through NCBI, are preceded by their NCBI accession. This tree was inferred from unaligned sequences using PASTA.

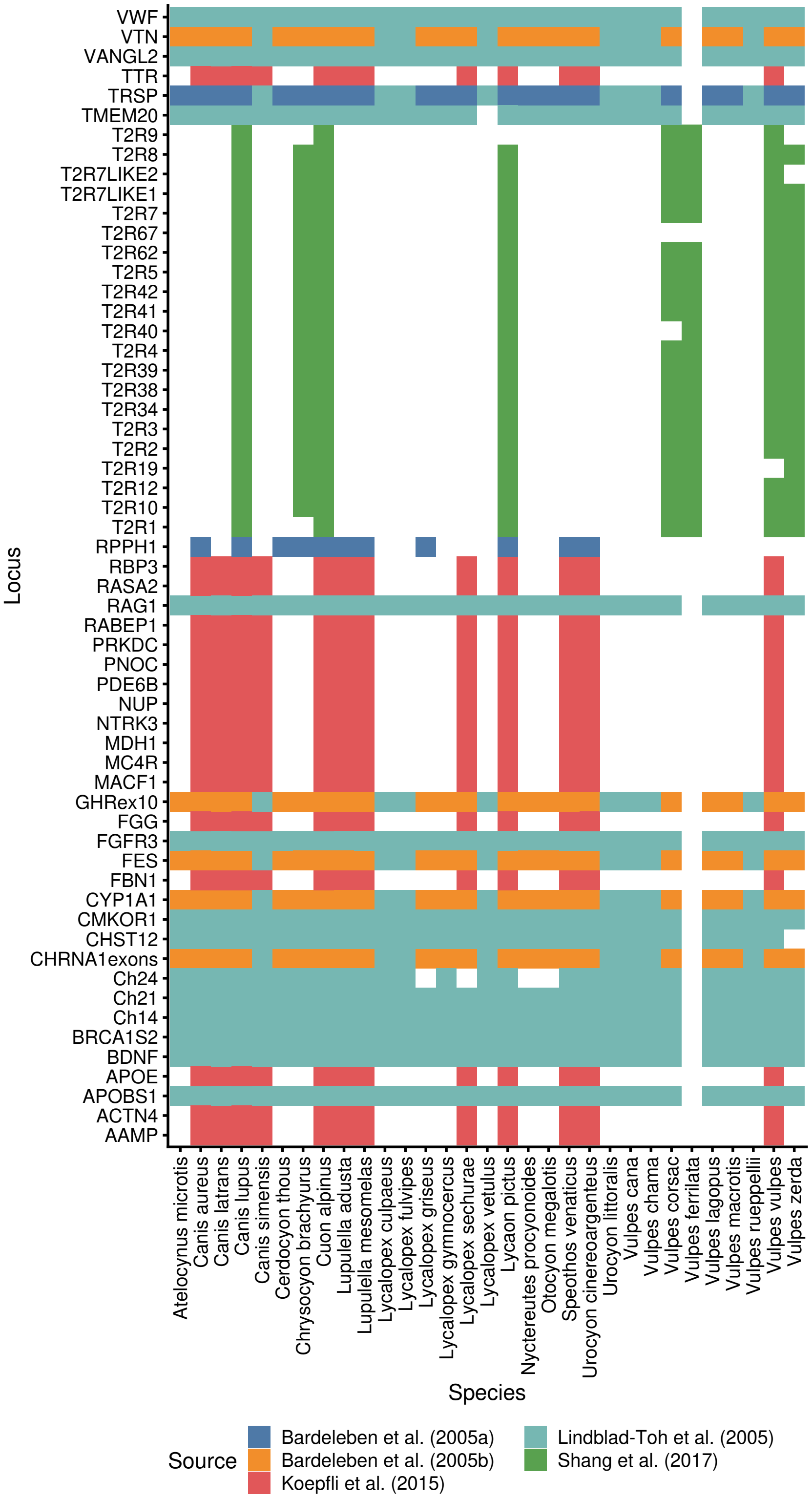


Figure S13: The loci and taxa in our molecular data set. Colored tiles show which of the five original sources [16, 28–31] a sequence was derived from, where blank tiles represent entirely missing sequences.

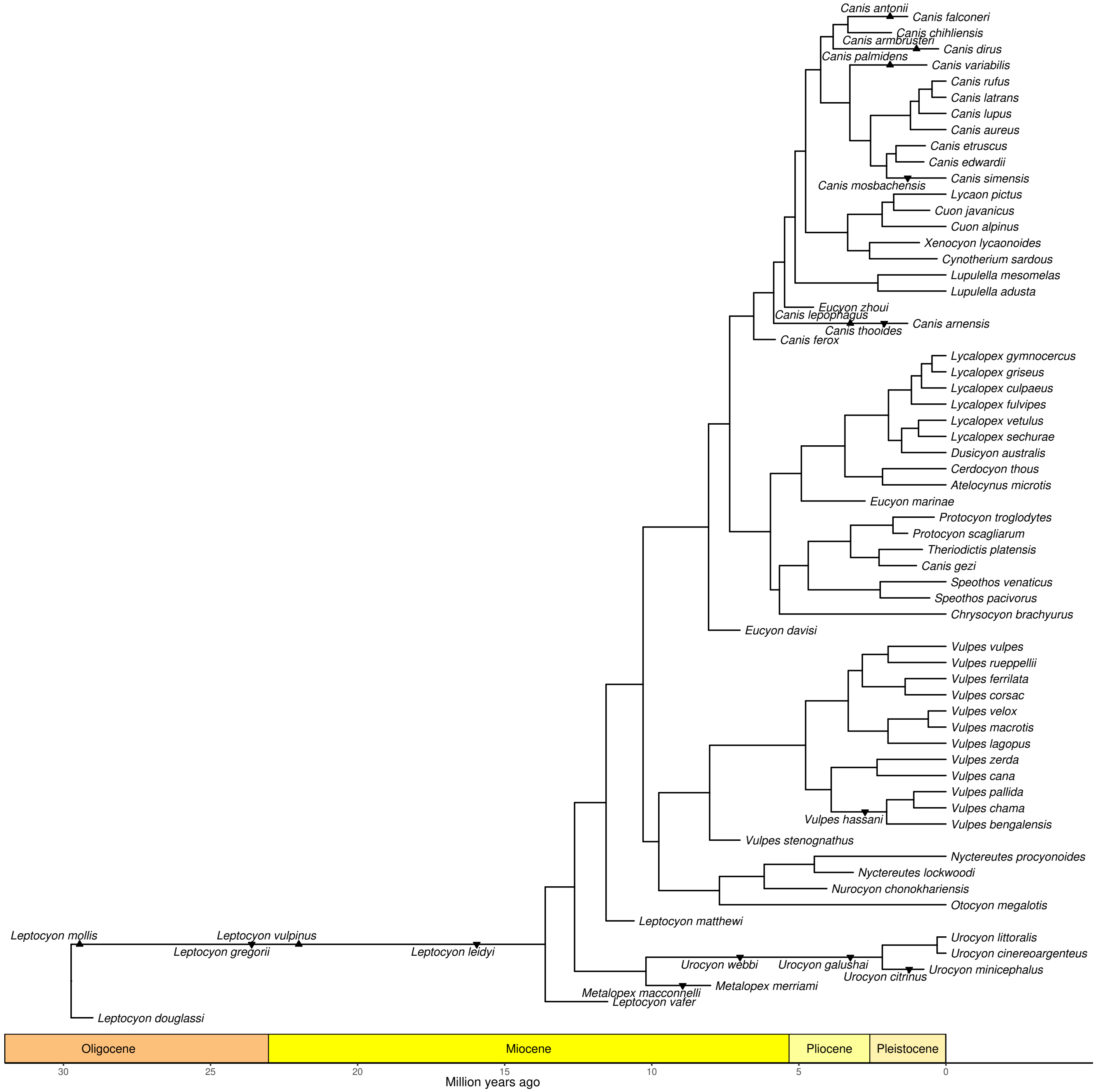


Figure S14: Maximum clade credibility tree summarizing FBD-MSC posterior distribution. Mean heights were used for node heights, and triangles represent sampled ancestors. Triangles point towards the species name for each sampled ancestor.

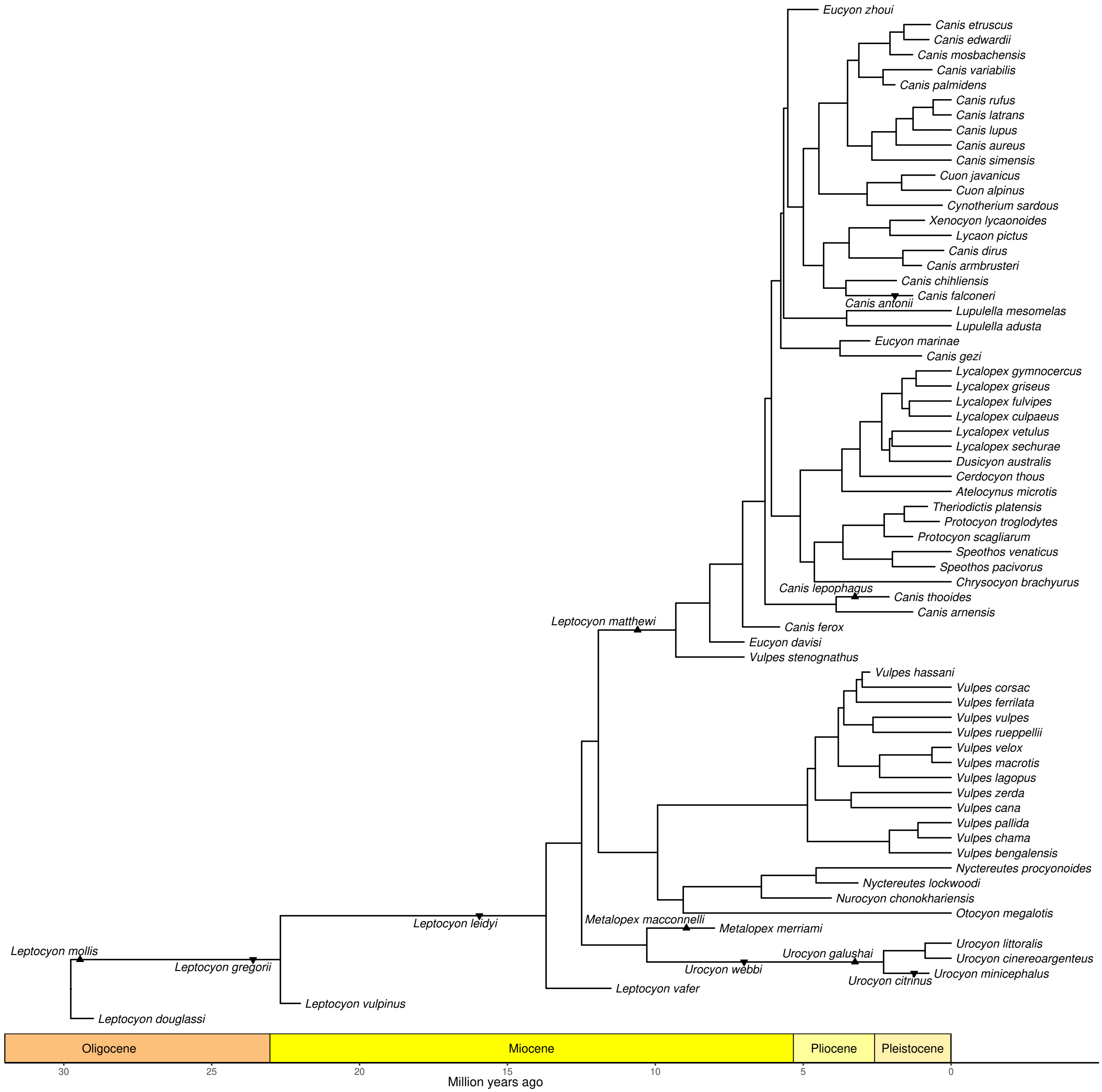


Figure S15: Maximum clade credibility tree summarizing FBD-concatenation posterior distribution. Mean heights were used for node heights, and triangles represent sampled ancestors. Triangles point towards the species name for each sampled ancestor.

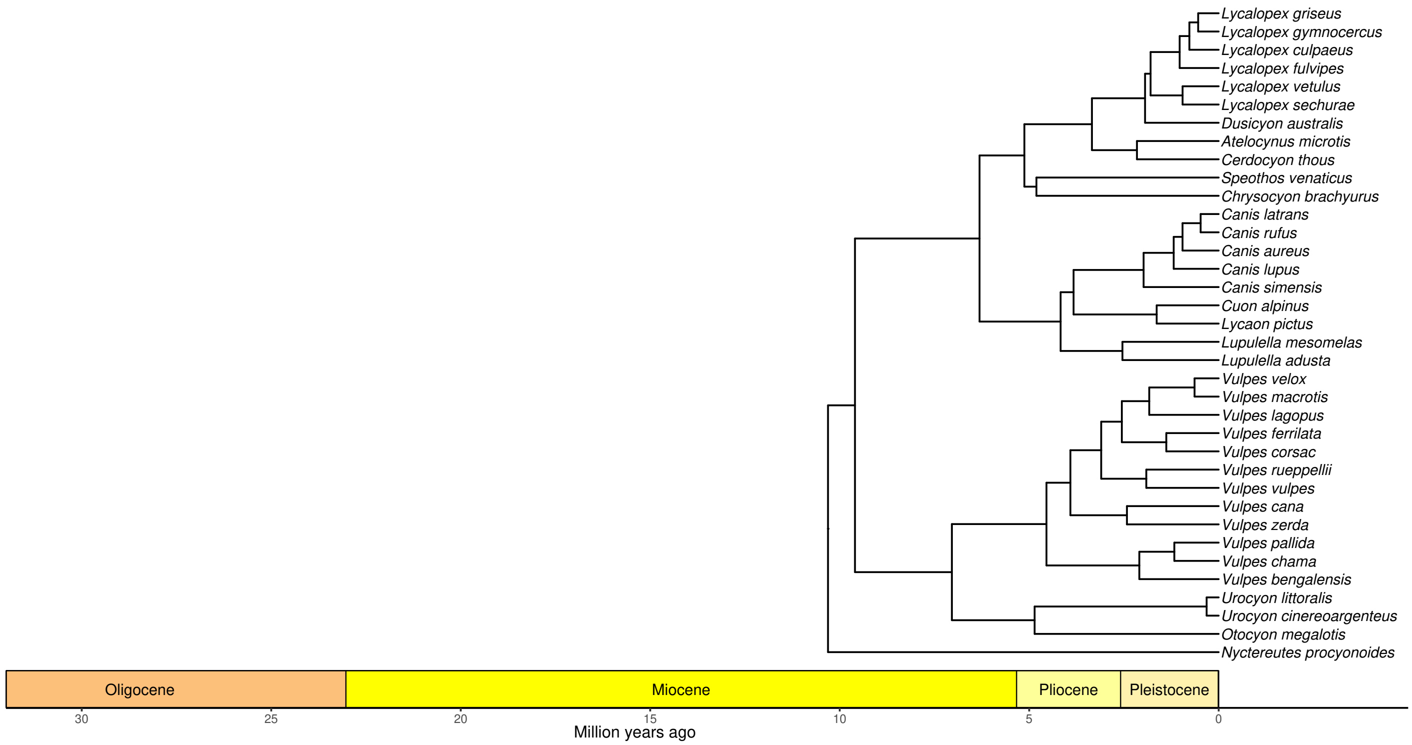


Figure S16: Maximum clade credibility tree summarizing BD-MSC posterior distribution. Mean heights were used for node heights.

### Additional tables

Table S7: Locus lengths and missing data.

| Locus | Length before[1] | Length after[2] | Missing[3] before[1] | Missing[3] after[2] | Total[4] missing before[1] | Total[4] missing after[2] |
|---|---|---|---|---|---|---|
| AAMP | 526 | 512 | 3.37% | 1.06% | 62.60% | 61.70% |
| ACTN4 | 529 | 501 | 4.54% | 0.00% | 63.05% | 61.29% |
| APOBS1 | 702 | 692 | 0.05% | 0.00% | 3.27% | 3.23% |
| APOE | 646 | 637 | 4.41% | 3.34% | 63.00% | 62.58% |
| BDNF | 489 | 486 | 0.61% | 0.43% | 3.81% | 3.64% |
| BRCA1S2 | 741 | 692 | 0.00% | 0.00% | 3.23% | 3.23% |
| Ch14 | 950 | 913 | 3.60% | 0.10% | 6.71% | 3.32% |
| Ch21 | 626 | 554 | 7.20% | 0.28% | 10.20% | 3.50% |
| Ch24 | 814 | 711 | 12.08% | 0.38% | 26.26% | 16.45% |
| CHRNA1exons | 383 | 314 | 6.48% | 0.00% | 9.49% | 3.23% |
| CHST12 | 705 | 695 | 0.00% | 0.00% | 6.45% | 6.45% |
| CMKOR1 | 735 | 735 | 0.00% | 0.00% | 3.23% | 3.23% |
| CYP1A1 | 619 | 590 | 4.97% | 1.73% | 8.04% | 4.90% |
| FBN1 | 683 | 673 | 1.31% | 0.24% | 61.80% | 61.38% |
| FES | 506 | 438 | 11.65% | 0.18% | 14.50% | 3.40% |
| FGFR3 | 503 | 487 | 2.22% | 0.01% | 5.37% | 3.23% |
| FGG | 678 | 664 | 1.88% | 0.25% | 62.02% | 61.39% |
| GHRex10 | 844 | 807 | 4.27% | 0.36% | 7.36% | 3.58% |
| MACF1 | 717 | 705 | 0.26% | 0.00% | 61.39% | 61.29% |
| MC4R | 857 | 842 | 0.10% | 0.00% | 61.33% | 61.29% |
| MDH1 | 604 | 558 | 7.91% | 7.41% | 64.35% | 64.16% |
| NTRK3 | 797 | 783 | 0.75% | 0.09% | 61.58% | 61.32% |
| NUP | 827 | 771 | 6.40% | 0.12% | 63.77% | 61.34% |
| PDE6B | 529 | 507 | 4.05% | 0.51% | 62.86% | 61.49% |
| PNOC | 259 | 252 | 0.00% | 0.00% | 61.29% | 61.29% |
| PRKDC | 756 | 742 | 1.52% | 0.26% | 61.88% | 61.39% |
| RABEP1 | 851 | 776 | 7.59% | 0.00% | 64.23% | 61.29% |
| RAG1 | 741 | 667 | 0.99% | 0.01% | 4.19% | 3.24% |
| RASA2 | 518 | 480 | 6.52% | 0.50% | 63.81% | 61.49% |
| RBP3 | 770 | 762 | 0.09% | 0.00% | 61.32% | 61.29% |
| RPPH1 | 698 | 669 | 9.61% | 6.32% | 67.93% | 66.76% |
| T2R1 | 894 | 894 | 0.00% | 0.00% | 77.42% | 77.42% |
| T2R2 | 912 | 912 | 0.00% | 0.00% | 74.19% | 74.19% |
| T2R3 | 951 | 951 | 0.00% | 0.00% | 74.19% | 74.19% |
| T2R4 | 900 | 897 | 0.04% | 0.00% | 74.20% | 74.19% |
| T2R5 | 888 | 888 | 0.00% | 0.00% | 74.19% | 74.19% |
| T2R7 | 945 | 936 | 0.60% | 0.00% | 74.35% | 74.19% |
| T2R7LIKE1 | 318 | 252 | 4.01% | 0.00% | 75.23% | 74.19% |
| T2R7LIKE2 | 932 | 367 | 51.96% | 0.08% | 89.15% | 77.44% |
| T2R8 | 1050 | 919 | 10.60% | 0.04% | 76.93% | 74.20% |
| T2R9 | 936 | 934 | 0.11% | 0.00% | 83.89% | 83.87% |
| T2R10 | 974 | 883 | 1.32% | 0.00% | 74.53% | 74.19% |
| T2R12 | 945 | 945 | 0.00% | 0.00% | 74.19% | 74.19% |
| T2R19 | 832 | 831 | 0.02% | 0.00% | 77.42% | 77.42% |
| T2R34 | 1014 | 807 | 10.54% | 0.00% | 76.91% | 74.19% |
| T2R38 | 951 | 942 | 0.12% | 0.00% | 74.22% | 74.19% |
| T2R39 | 963 | 963 | 0.00% | 0.00% | 74.19% | 74.19% |
| T2R40 | 921 | 915 | 0.09% | 0.00% | 77.44% | 77.42% |
| T2R41 | 927 | 927 | 0.00% | 0.00% | 74.19% | 74.19% |
| T2R42 | 974 | 970 | 0.21% | 0.00% | 74.25% | 74.19% |
| T2R62 | 1211 | 885 | 23.41% | 2.54% | 80.23% | 74.85% |
| T2R67 | 944 | 937 | 0.79% | 0.28% | 80.80% | 80.70% |
| TMEM20 | 615 | 596 | 1.05% | 0.12% | 7.44% | 6.56% |
| TRSP | 2167 | 704 | 67.43% | 0.56% | 68.48% | 3.77% |
| TTR | 1070 | 1028 | 1.08% | 0.00% | 61.71% | 61.29% |
| VANGL2 | 546 | 520 | 0.16% | 0.00% | 3.38% | 3.23% |
| VTN | 487 | 481 | 2.43% | 1.40% | 5.58% | 4.58% |
| VWF | 732 | 721 | 0.05% | 0.00% | 3.27% | 3.23% |
| Concatenated | 45602 | 41620 | 9.50% | 0.41% | 54.78% | 51.56% |

[1] Before trimming
[2] After trimming
[3] Percentage missing data among species with a sequence for a given locus
[4] Percentage missing data including species without sequences for a given locus (but which have sequences for other loci)

Table S8: Effective population sizes in canid species. We assume a generation time of 4 years [32], and report $N_e$ bounds given $\frac{N_e}{N}$ is 0.1 and 0.8 [33].

| Species name | Population size estimate | $N_e$ | Reference |
|---|---|---|---|
| *Urocyon littoralis* | 4,000 | 400 – 3200 | [34] |
| *Lycaon pictus* | 6,600 | 660 – 5,280 | [35] |
| *Chrysocyon brachyurus* | 17,000 | 1700 – 13,600 | [36] |
| *Speothos venaticus* | 110,000 | 11,000 – 88,000 | [37] |
| *Canis lupus* | 200,000–250,000 | 20,000 – 200,000 | [38] |
| *Canis latrans* | $> 113,000$ | >11,300 – >90,400 | [39] |
| *Vulpes lagopus* | $> 200,000$ | >20,000 – >160,000 | [40] |
| *Vulpes vulpes* | 509,000–7,275,000 (UK) | >72,000 – >5,760,000 | [41] |
